# Paclitaxel Dosing Regimens Drive Differential CD4⁺ T Cell Responses in the Dorsal Root Ganglia and Modulate Neuropathic Pain Severity

**DOI:** 10.64898/2026.06.05.730418

**Authors:** Kenyah G.N. Ferreira, Moriah E. Weese-Myers, Leslie S. Bradford, Diana J. Goode

## Abstract

While chemotherapy is effective in killing cancer cells, its non-specific cytotoxicity harms peripheral neurons, leading to chemotherapy-induced peripheral neuropathy (CIPN). Lacking effective interventions, clinicians often reduce or discontinue treatment. However, these adjustments do not necessarily reverse neuropathy and may jeopardize cancer control. Growing evidence now indicates that chemotherapy not only exerts neurotoxic effects but also modulates immune responses. The degree to which chemotherapy dosing regimens shape CD4^+^ T cell responses in the dorsal root ganglia (DRG), and how these cells influence mechanical hypersensitivity, remains poorly understood. In this study, female mice were administered a single, high dose (sHD) or multiple low doses (mLD) of paclitaxel (PTX). The mLD group, which received the higher cumulative dose, exhibited an earlier T cell response in the DRG and attenuated mechanical hypersensitivity with fewer ATF3^+^ DRG neurons compared to the sHD group. This regimen promoted a focused CD4^+^ T cell response while driving a broad and diversified CD8^+^ T cell expansion. In contrast, the sHD PTX regimen elicited a delayed, polyfunctional CD4^+^ T cell response but generated limited CD8^+^ effector differentiation. To directly assess the contribution of CD4^+^ T cells to CIPN pathogenesis, we administered PTX to mice lacking CD4^+^ T cells. CD4 deficient mice were significantly less hypersensitive than CD4^+^ sufficient mice with a stronger reduction in the sHD group. Although higher cumulative doses are associated with increased CIPN risk, our results suggest that the concentration and frequency of PTX more directly influence DRG immune programming, and that this immune shaping modulates CIPN severity.

## INTRODUCTION

Chemotherapy-induced peripheral neuropathy (CIPN) is a common and debilitating adverse effect of cancer treatment, particularly with paclitaxel (PTX). Patients frequently develop numbness, tingling, and mechanical and cold hypersensitivity in a stocking–glove distribution. These sensory abnormalities are associated with pathological changes in dorsal root ganglion (DRG) neurons, including hyperexcitability, axonal toxicity, and neuroinflammation, which together drive neuropathic pain. There are no consistently effective treatments for CIPN; as a result, chemotherapy doses are often reduced, delayed, or discontinued. However, a recent study reported that dose modification did not necessarily improve neuropathy outcomes [62], highlighting the clinical challenge of balancing toxicity and efficacy. Importantly, ∼26% of patients continue to experience neuropathy long after completing treatment[56].

Mounting evidence indicates that CIPN arises from both neuronal injury and immune-mediated mechanisms. PTX induces inflammatory signaling within the DRG, where infiltrating macrophages release pro-inflammatory cytokines such as TNF-α that enhance neuronal excitability[69]. In addition to innate immune activation, adaptive immune cells—particularly T lymphocytes—play critical roles in modulating neuropathy. Rag^−/-^ mice (lack adaptive cells) exhibit prolonged mechanical hypersensitivity after PTX, and intravenous transfer of CD8⁺ T cells promotes resolution[26]. Conversely, intrathecal transfer of effector CD4⁺ or CD8⁺ T cells exacerbates hypersensitivity, whereas transfer of regulatory T cells (Tregs) reduces pain behaviors[33]. Neuron-specific reduction of MHCII decreases anti-inflammatory CD4⁺ T cells in the DRG and worsens PTX-induced cold hypersensitivity[65], underscoring bidirectional neuron–T cell interactions. PTX induces both pro- and anti-inflammatory CD4⁺ T cells in the DRG[13], and anti-inflammatory cytokines such as IL-10[26] and IL-4[55] are protective. Together, these studies suggest that pro-inflammatory T cells promote CIPN, whereas anti-inflammatory T cells suppress its development and facilitate resolution.

CIPN is often described as dose-dependent and cumulative, yet preclinical findings reveal important complexity. Higher cumulative dosing (8 mg/kg) produces prolonged hypersensitivity lasting >90 days[63]. However, lower regimens (2 and 4 mg/kg) do not differ in magnitude or recovery time[36; 63]. Beyond behavioral effects, PTX exerts dose-dependent immunomodulation. At ultra-low doses (1 mg/kg), PTX reduces Tregs and myeloid-derived suppressor cells (MDSCs) while enhancing IFN-γ and effector T cell responses[14; 28]. Low doses (4 mg/kg) do not markedly suppress T cell numbers, whereas moderate to high doses reduce major T cell subsets and alter regulatory balance[15]. PTX concentration also influences dendritic cell (DC) maturation and antigen presentation[44], shifting from immunostimulatory at low doses to immunosuppressive at higher concentrations[19]. In parallel, dosing schedule shapes immune programming: single high doses cause transient inflammation, metronomic regimens often preserve effector responses while suppressing regulatory populations[7], and repeated cycles can drive immune dysregulation[52].

Despite these advances, prior CIPN studies have used diverse PTX dosing regimens (1–8 mg/kg; variable injection schedules), limiting direct comparisons of immune outcomes. A key knowledge gap remains: how do PTX dose intensity and administration schedule differentially modulate CD4⁺ T cell subsets within the DRG, and how do these immune shifts influence the magnitude and persistence of mechanical hypersensitivity? Addressing this question will clarify how chemotherapy regimens shape neuroimmune interactions and may inform strategies to mitigate CIPN without compromising anti-tumor efficacy.

## MATERIALS AND METHODS

### Mice

This study was conducted exclusively in female mice to allow for sufficient statistical power and depth of analysis within a single sex. Given the recognized importance of sex as a biological variable, complementary studies in male mice are ongoing and will be the focus of a subsequent report. Female C57BL6/J mice were purchased from The Jackson Laboratory. Mice were given food and water ad libitum. All experimental protocols followed National Institutes of Health guidelines and were approved by the University of New England Institutional Animal Use and Care Committee. All experiments were performed with 6–12-week-old female C57BL6/J mice. Mice were euthanized with an overdose of avertin (0.5 cc of 20 mg/ml) followed by transcardiac perfusion with ice-cold 1× PBS (137 mM NaCl, 2.7 mM KCl, 10 mM Na_2_HPO_4_, and 1.8 mM KH_2_PO_4_, pH = 7.40 ± 0.01). This method of euthanasia is consistent with American Veterinary Medical Association (AVMA) Guidelines for the Euthanasia of Animals.

### Paclitaxel dosing regimens

Paclitaxel (PTX, Sigma-Aldrich, T7191) was solubilized in cremophor:ethanol 1:1 mixture and diluted 1:3 in sterile saline. Two PTX dosing regimens were used: a single high dose of 6 mg/kg PTX administered on day 0 (sHD, as done previously[13; 33; 65]) or multiple low doses of 2 mg/kg PTX administered on days 0, 2, 4, 6 (mLD, cumulative dose 8 mg/kg, as done previously[10; 35]) **(Figs 1A, S1)**. Although these PTX regimens are frequently used in prior studies, our selection was guided by immunological considerations related to T cell activation and differentiation. One regimen was specifically designed to produce a higher cumulative dose, as greater PTX exposure is widely believed to correlate with increased CIPN severity. PTX for both regimens were given as an intraperitoneal injection (IP) at a volume of 10 ml/kg bodyweight. For the CD4^+^ T cell deficient (GK1.5, BioXcell, BE0003-1) and sufficient (LTF.2 isotype control, BioXcell, BE0090) experiments, antibodies were prepared at a concentration of 9 mg/kg (diluted in InVivoPure Buffers from BioXcell: pH 6.5 for GK1.5 and pH 7.0 for LTF.2) and administered as an IP injection on days -4, -3, 4, 8, 15, and 22 **(S1 Fig)**. Mice were randomly assigned a treatment, which experimenters were blinded to during testing.

**Figure 1.**
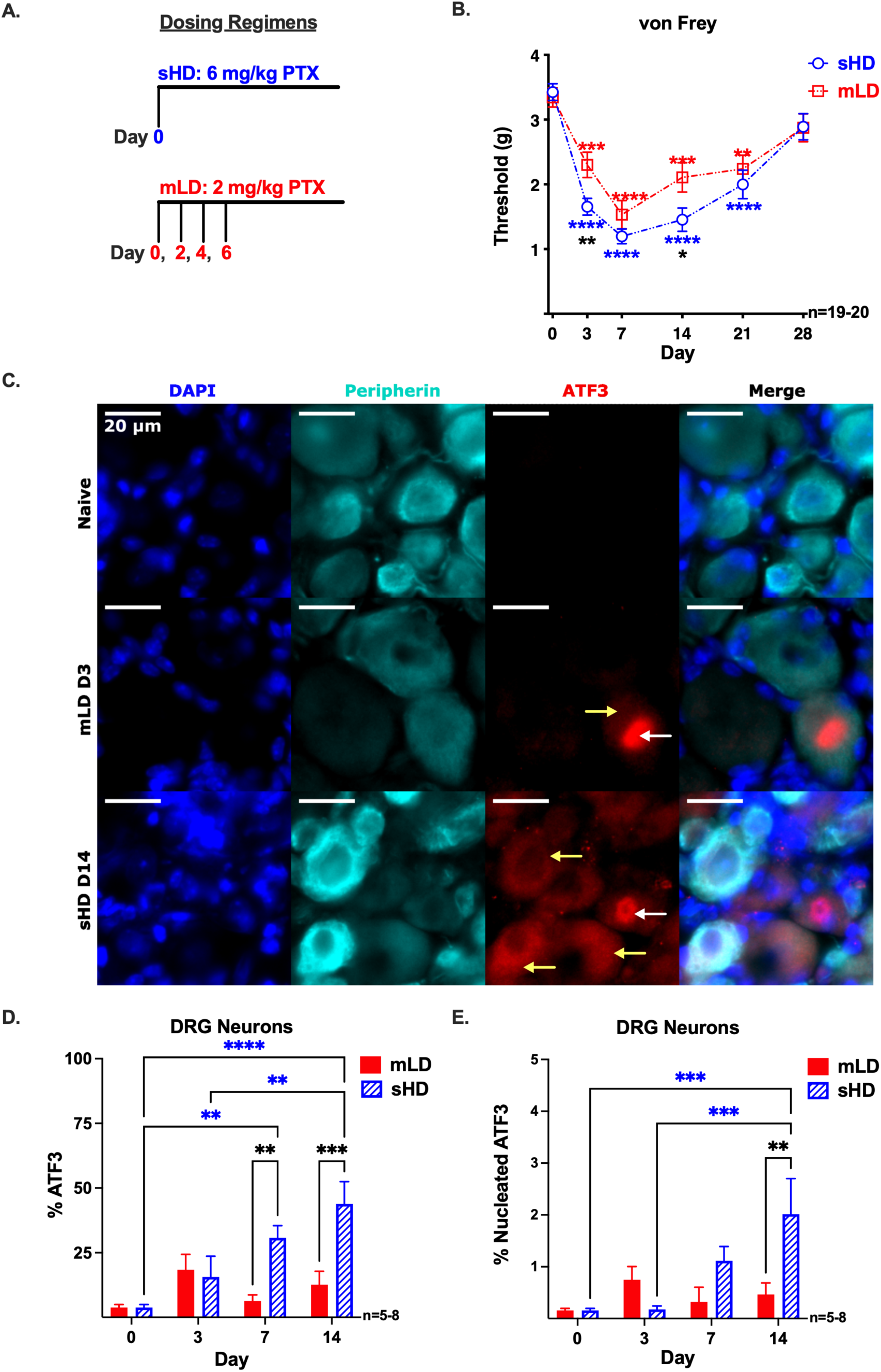
The sHD PTX regimen induces more severe mechanical hypersensitivity and the activation of neuronal ATF3 than the mLD regimen. **(A)** Female C57BL6/J mice received either a single high dose of 6 mg/kg PTX administered on day (D) 0 (sHD, blue) or multiple low doses of 2 mg/kg PTX given on D 0, 2, 4, 6 (mLD, red). **(B)** Mechanical hypersensitivity (tactile threshold in grams) was assessed with von Frey filaments prior to PTX on D 0 to determine baseline (BL) values and after sHD or mLD PTX on D 3, 7, 14, 21, and 28. Significance was determined by 2-way repeated measures ANOVA with Tukey’s multiple comparisons test (*p<0.05, **p<0.01, ***p<0.001, ****p<0.0001, n=20/regimen). Blue (sHD) and red (mLD) significance asterisks compare hypersensitivity after PTX to BL within each dosing regimen. Black significance asterisks compare hypersensitivity values between sHD and mLD PTX regimens. **(C)** Representative widefield epifluorescence images of peripherin (teal) L4 DRG neurons from naïve, mLD PTX (day **(D)** 3), and sHD PTX (day (D) 14) female mice. Sub-cellular location of ATF3 (red) is indicated by arrows: yellow denotes cytoplasmic expression, and white denotes nuclear localization (co-localized with DAPI, blue). Quantification of (D) the percentage of ATF3^+^ peripherin^+^ neurons, and **(E)** the percentage of peripherin^+^ neurons with nuclear localization of ATF3. **(D, E)** Statistical significance determined by 2-way ANOVA with Tukey’s multiple comparison test (**p<0.01, ***p<0.001, ****p<0.0001, n=5-8). All data presented as mean ± SEM.

### Immunohistochemistry (IHC)

Mice were euthanized with avertin and transcardially perfused with ice-cold 1× PBS followed by ice-cold 4% paraformaldehyde (PFA). Lumbar 4 (L4) DRGs were harvested and post-fixed in 4% PFA for 1 hour at room temperature (RT), then cryoprotected in 30% sucrose containing 0.02% sodium azide at 4 °C for up to one week. DRGs were embedded in Clear Frozen Section Compound (VWR), stored at −80 °C, and later serially sectioned at 12 μm. Prior to staining, slides were equilibrated to RT, and the embedding compound was removed via three 5-minute washes in 1× PBS. Tissue was permeabilized with 0.1% Triton-X in PBS (PBS-T) for 15 minutes at RT, followed by a 1-hour block in 5% normal donkey serum (NDS). Slides were incubated overnight at RT in a light-protected, humidified chamber with primary antibodies diluted in NDS-PBS-T (Peripherin, Invitrogen, PA5-21031; ATF3, Novus Biologicals, NBP1-85816). After three 5-minute washes, slides were incubated with secondary antibodies (Donkey anti-chicken 488, Jackson ImmunoResearch, 703-165-155; Donkey anti-rabbit Cy3, Jackson ImmunoResearch, 711-545-152) for 1 hour at RT. Slides were washed again, then coverslipped using Fluoroshield Mounting Medium with DAPI (Abcam). Slides were imaged on the day of staining using a Keyence BZ-X710 automated widefield fluorescent microscope. Images were acquired with a 40X objective (Plan Fluor, NA 0.75) using a cooled monochrome CCD camera (960×720 resolution, 14-bit gradation).

### IHC automated immunofluorescent image analysis

ATF3^+^ and nucleated ATF3^+^ DRG neurons were quantified using a custom automated module in the MetaXpress software (Molecular Devices). Before analysis, multi-channel images were split into single-channel images in Fiji. The quantification pipeline was defined by four steps:

1. Neuron Cell Body Identification: A mask for neuronal cell bodies was generated based on peripherin intensity and cell size, specifically excluding nerve fibers.
2. Threshold Calibration: The threshold for ATF3 positivity was set using the mean intensity of nerve fiber background staining from naïve mouse DRGs. This conservative baseline was used because the isotype control (99th percentile) intensity was lower than the intrinsic background of the nerve fibers.
3. Nuclear Masking: A DAPI-based mask was used to identify nuclei.
4. Logical Classification: Nucleated ATF3^+^ neurons were identified using a logical operation requiring the overlap of the nuclear and cell body masks, with an ATF3 mean intensity ≥ 5000 MPI above local background.

To ensure statistical robustness, the automated module evaluated an average of 1330 ± 105 neurons per DRG. Representative images were selected from the mouse whose percent of ATF3^+^ neuron population most closely matched the group mean for each dosing regimen and timepoint. Images were processed in Fiji as 16-bit files (0–65,535 MPI). To maintain consistency between the visual figures and the quantitative analysis, background staining below the analysis threshold for ATF3 (5,000 MPI) was removed. For visualization purposes, an upper display threshold of 20,000 MPI was applied to the ATF3 channel to enhance signal contrast. Images were pseudocolored to clearly distinguish individual markers.

### Assessment of mechanical sensitivity (von Frey test)

von Frey was used to quantify evoked measures of mechanical hypersensitivity in naïve female mice and PTX-treated on days -5, 0, 3, 7, 14, 21, and 28 **(S1 Fig)**. In this test, mice were placed on a mesh screen within a plexiglass chamber. A von Frey monofilament of 3.61, which is equivalent to 0.41 g force, was applied to the plantar surface of the hindpaw with enough force to bend the filament for 2–3 s. Paw withdrawal during and immediately after removal of the stimulus was considered a positive response and recorded as “X” in the recording template. A positive response resulted in the use of the next lowest filament in the series (3.22). A negative response (recorded as “O” in the recording template) to the initial 3.61 filament resulted in the use of the next highest filament. Testing of higher filaments were used until a positive response was reached (4.08, 4.31, 4.56). This up/down method as previously reported[12] was continued until 4 filaments were tested after the initial positive response. Mechanical hypersensitivity was determined by entering the response pattern in the up-down Excel program generously provided by Dr. Michael Ossipov (University of Arizona, Tucson, AZ).

### DRG cell isolation and flow cytometry

DRGs (60) were collected and pooled for each mouse. DRGs were acutely dissociated as described previously[65]. Single cell suspensions of dissociated DRGs were passed through a 70 μM cell strainer to eliminate clumps and tissue debris. Cells were incubated at room temperature for 30 minutes with Live/Dead Fixable Violet (ThermoFisher) and 1× brefeldin A solution (BioLegend) for the intracellular cytokine panel **(Supplemental Table 1)**. Cells were incubated with anti-mouse CD16/32 (Biolegend: TruStain FcXTM) for 10 minutes prior to the addition of extracellular antibodies. **Supplemental Tables 1** and **2** list the antibodies used for the cytokine/transcription factor panel **(Figs 2, 3, S3, 4)** and the immune cell population panel **(Figs 5B, 5E-F, S5B)**, respectively. Cell surface antibodies were added to the cells for 20 minutes at 4 °C. Cells were washed with FACs buffer (1× PBS, 1% FBS) and fixed at room temperature for 20 minutes with Intracellular Staining Fixation Buffer (BioLegend). Cells were washed with 1× Cyto-FastTM Perm Wash solution (BioLegend) and incubated with intracellular antibodies for 20 minutes at room temperature. VersaComp Antibody Capture Kit (Beckman Coulter) was used to set compensation to correct for spectral overlap. Unstained DRG cells were used to check for auto-fluorescence. Isotype controls were used to determine non-specific binding and to set cell population gates. Gating strategies are shown in **(S2 Fig)**. Greater than 100,000 events in the live DRG cell gate were acquired on the Beckman Coulter CytoFLEX S System B2-R3-V4-Y4 and data analyzed with FlowJo^TM^ 10.9.0.

**Figure 2.**
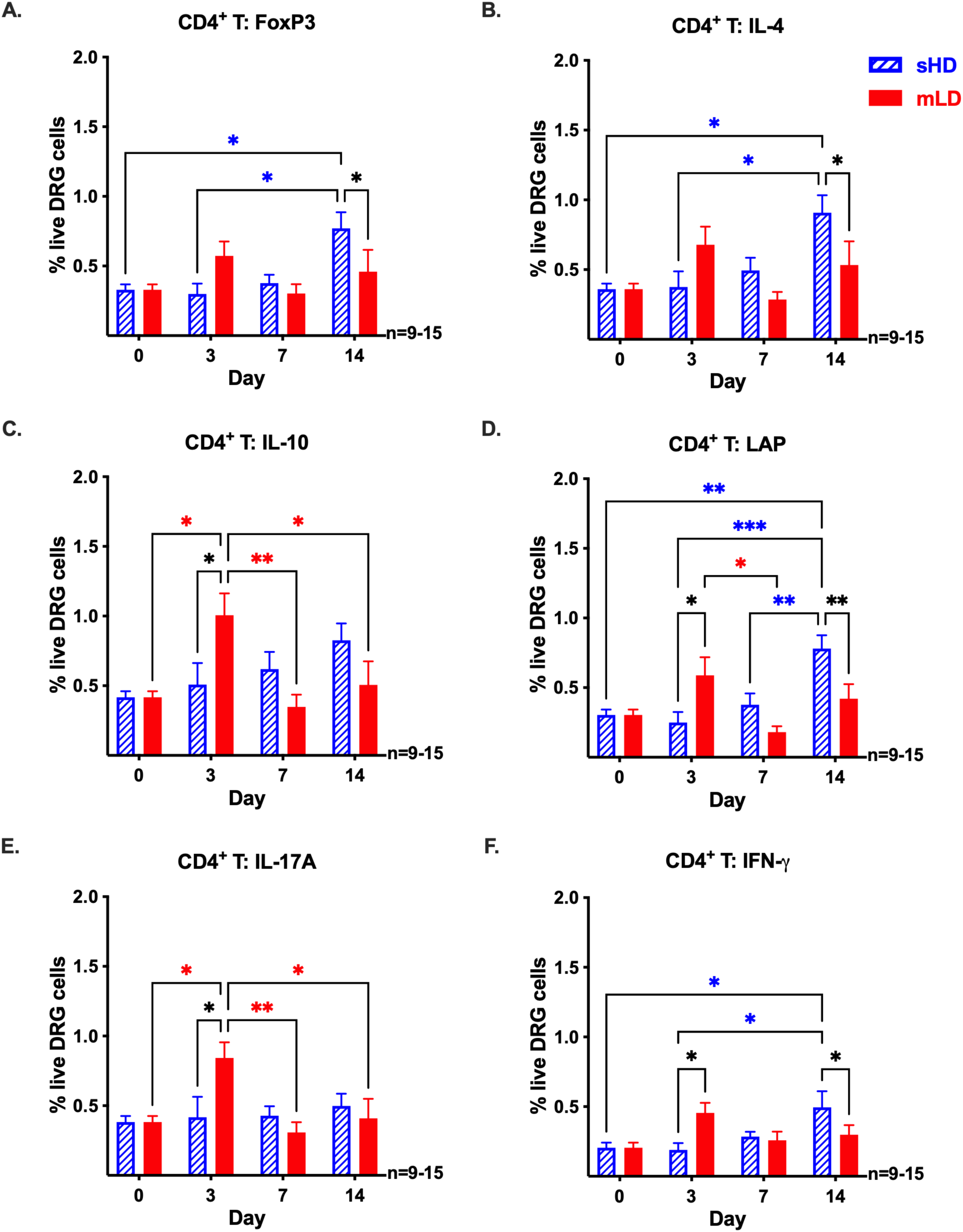
The mLD PTX regimen induces a faster but more restricted CD4^+^ T cell response in the DRG than the sHD regimen. Frequencies of **(A)** FoxP3, **(B)** IL-4, **(C)** IL-10, **(D)** LAP, **(E)** IL-17A, and **(F)** IFN-γ CD4^+^ T cells out of total live DRG cells on day 0 and after sHD (blue) or mLD (red) PTX regimens. Significance was determined by 2-way ANOVA with Tukey’s multiple comparisons test (*p<0.05, **p<0.01, ***p<0.001, n=9-15/regimen). Blue (sHD) and red (mLD) significance asterisks compare CD4^+^ T cell frequencies within each dosing regimen. Black significance asterisks compare CD4^+^ T cell frequencies between sHD and mLD PTX regimens.

### Statistical analysis

Sample sizes were determined based on historical data from our laboratory to ensure sufficient sensitivity for detecting medium-to-large effect sizes. To assess the magnitude of the observed biological changes independently of sample size, we calculated partial eta squared (ηp2) for all ANOVA main effects and interactions. Effect sizes were interpreted according to Cohen’s criteria, where 0.01, 0.06, and 0.14 represent small, medium, and large effects, respectively (**Supplemental Table 3**). All experiments were analyzed using Graphpad Prism 10 (Graphpad Software, Inc). Behavior data were checked for normality (Shapiro-Wilk), equal variances (F test), and outliers (ROUT). Two-way repeated measures ANOVA with Tukey’s multiple comparisons tests were used to determine significance in the von Frey behavior tests **(Figs 1B, 5C, 5D, S5, S6A)**. A two-way ANOVA with Tukey’s multiple comparisons tests was used to compare the mean percentage of ATF3^+^ and nucleated ATF3^+^ DRG neurons between PTX dosing regimens and time. Normality was tested via QQ plot and Shapiro-Wilk **(Figs 1D, 1E)**. Flow cytometry data **(Figs 2, 3, 4, S4)** were analyzed by a two-way ANOVA with Tukey’s multiple comparisons test. A one-way ANOVA was used to compare the mean frequency of neutrophils in the DRG of naïve (± LTF.2) and day 14 PTX treated mice (sHD and mLD ± LTF.2) **(S6B)**. A one-way ANOVA was used to compare the mean frequency of cytokine producing immune cells (CD3^−^, CD8^+^ T cells) in the DRG of naïve and CD4^+^ T cell sufficient (LTF.2) and deficient (GK1.5) day 14 PTX (sHD and mLD) treated mice **(Figs 6A-D, 7A-D, S7)**. Immune cell population frequencies are presented as fold-changes between CD4^+^ T cell sufficient mice (LTF.2) compared to CD4^+^ T cell depleted mice (GK1.5); however, statistical significance was determined by an unpaired t-test with Welch’s correction using immune cell frequencies (not on the fold-change) between CD4^+^ T cell sufficient mice (LTF.2) and CD4^+^ T cell depleted mice (GK1.5) within each cell subset **(Fig 5B, E, F)**. Results are reported as mean ± SEM with p values <0.05 to be considered significant. Degrees of freedom, F and p scores, and effect sizes are presented in **Supplemental Table 3**. Raw data, mean, SEM, and 95% confidence intervals are presented in **Supplemental Table 4**.

## RESULTS

### sHD PTX regimen induces greater mechanical hypersensitivity and activates neuronal ATF3 expression

A higher cumulative dose of chemotherapy is often recognized as a significant risk factor for CIPN[1; 67]; however, many studies suggest that this relationship is complex with many other contributing factors. To investigate this further, we designed two dosing regimens – a single high dose of 6 mg/kg PTX administered on day 0 (sHD), and a multiple low dose regimen of 2 mg/kg PTX given on days 0, 2, 4, and 6 (mLD) **(Fig 1A)**. On day 3, both the mLD and sHD mice were significantly more hypersensitive to mechanical stimulation compared to baseline (day 0) values assessed by von Frey (mLD: 2.302 g ± 0.196 SEM, p=0.0003, sHD: 1.656 ± 0.131, p<0.0001; n=19-20) with the sHD group significantly more hypersensitive than the mLD group (p=0.0099) **(Fig 1B)**. Thus, at day 3, the mice that received the higher cumulative dose (sHD: 6 mg/kg) were more hypersensitive than the mice that received the lower cumulative dose (mLD: 4 mg/kg). On day 7, both groups reached peak hypersensitivity to the same severity (mLD: 1.533 ± 0.219, p<0.0001 and sHD: 1.196 ± 0.113, p<0.0001, compared to baseline values) even though the mLD group had a higher cumulative dose (8 mg/kg) compared to the sHD group (6 mg/kg). By day 14, the PTX-induced mechanical hypersensitivity started to resolve (mLD: 2.107 ± 0.227, p=0.0014 and sHD: 1.456 ± 0.182, p<0.0001, compared to baseline values); however, the mLD group resolved faster than the sHD group (p=0.0318), even though the mLD group had a higher cumulative dose. By day 21, both groups continued to resolve (mLD: 2.239 ± 0.212, p=0.0027 and sHD: 1.999 ± 0.221, p<0.0001, compared to baseline values) with no difference in the severity of mechanical hypersensitivity. By day 28, both groups had returned to baseline values. These results suggest that other factors induced by PTX dosing (concentration and frequency) contribute to mechanical hypersensitivity.

While mechanical hypersensitivity can occur without overt neuronal damage[5], PTX has been shown to increase expression of Activating Transcription Factor 3 (ATF3)[31], a marker for neurons in an injury-associated transcriptional state. Since PTX preferentially affects c-fibers and Aδ-fibers[45], we investigated the extent to which the two PTX dosing regimens induced ATF3 in peripherin^+^ nociceptive neurons in L4 DRG tissue **(Fig 1C-E, S3 Fig)**. In naïve mice **(Fig 1C top row)**, ATF3 expression was minimal (3.82% ± 1.141) **(Fig 1D)** with almost no nuclear localization (0.158% ± 0.037) **(Fig 1E)**. Following treatment with mLD PTX **(Fig 1C middle row, S3 Fig)**, there was a non-significant increase in ATF3^+^ peripherin^+^ neurons (3.82% ± 1.141 in naïve to 18.44% ± 5.941 at day 3, p=0.2162, n=7-8) **(Fig 1C)** and peripherin^+^ neurons where ATF3 localized to the nucleus (0.158% ± 0.037 in naïve to 0.750% ± 0.254 at day 3, p=0.5626, n=7-8) **(Fig 1D)**. In contrast to the mLD group, the sHD PTX regimen induced a robust and significant increase in ATF3 expression **(Fig 1C bottom row, S3 Fig)**. By day 7, 30.77% ± 4.71 of peripherin^+^ neurons were ATF3^+^ compared to naïve (3.82% ± 1.141, p=0.0026, n=7-8), a response significantly greater than that seen in the mLD group at day 7 (6.35% ± 2.333, p=0.0014, n=8). This effect was even more pronounced by day 14, reaching 43.84% ± 8.64 in the sHD group compared to naïve (3.82% ± 1.141, p<0.0001, n=8), which was markedly higher than the mLD group at the same timepoint (12.63% ± 5.21, p=0.0004, n=5-8) **(Fig 1D)**. Furthermore, the sHD regimen significantly increased the nuclear localization of ATF3 by day 14 (2.02% ± 0.69), representing a significant increase over both naïve (0.466% ± 0.222, p=0.0005, n=5-8) and the mLD group (0.47% ± 0.22, p=0.0031, n=5-8) **(Fig 1E)**. Collectively, these data demonstrate that the sHD PTX regimen reaches a critical pathological threshold, simultaneously driving significant mechanical hypersensitivity and activating the ATF3-mediated neuronal stress pathway.

### mLD PTX regimen induces a faster but restricted CD4^+^ T cell response in the DRG compared to sHD PTX

One potential reason for reduced CIPN in the mLD group is that this dosing regimen may induce different CD4^+^ T cell responses (breadth and depth) as both the intensity and duration of T cell stimulation result in unique subtypes of T cells[20]. Therefore, we characterized the anti- and pro-inflammatory CD4^+^ T cells in the DRG at baseline (day 0) and after administration of the two PTX dosing regimens on days 3, 7, and 14. Like we have found previously[13], the sHD PTX regimen induced a robust anti-inflammatory FoxP3 (>2.3-fold: 0.33% ± 0.038 of total live DRG cells at baseline to 0.77% ± 0.115 at day 14, p=0.0254, n=9-13) **(Fig 2A)** and IL-4 (>2.5-fold: 0.36% ± 0.040 of total live DRG cells at baseline to 0.91% ± 0.125 at day 14, p=0.0114, n=9-13) **(Fig 2B)** CD4^+^ T cell response at day 14. Furthermore, the sHD regimen induced significantly higher frequencies of IL-4 CD4^+^ T cells (0.91% ± 0.125 of total live DRG cells for sHD compared to 0.53% ± 0.170 for the mLD, p=0.0150, p=9-15) than the mLD group. In contrast, the mLD regimen induced a robust anti-inflammatory IL-10 CD4^+^ T cell response in the DRG at day 3 (>2.5-fold: 0.42% ± 0.043 of total live DRG cells at baseline to 1.01% ± 0.158 at day 3, p=0.0240, n=9-15) **(Fig 2C)**, which was significantly greater than the frequency for the sHD mice (0.51% ± 0.155, p=0.0162, p=9). Thus, the mLD group, which had a faster anti-inflammatory CD4^+^ T cell response in the DRG, also had reduced PTX-induced mechanical hypersensitivity.

Most often, when tissue injury occurs, both pro- and anti-inflammatory immune cell subtypes are induced as pro-inflammatory cells respond to signals of threats while anti-inflammatory cells prevent excessive tissue damage[6; 59]. Moreover, some cytokines like transforming growth factor-beta (TGF-β) are pleiotropic, meaning it can either promote pro- or anti-inflammatory responses depending on other cytokines present in the local microenvironment[29]. To detect cell-associated TGF-β, we used an antibody against latency associated peptide (LAP), the pro-peptide of TGF-β, as the epitope of mature TGF-β is physically masked by LAP. We found that the sHD induced a robust increase in LAP^+^ CD4^+^ T cells in the DRG at day 14 (>2.5-fold: 0.30% ± 0.039 of total live DRG cells at baseline to 0.78% ± 0.095 at day 14, p=0.0016, n=9-13) **(Fig 2D)**, which was greater than the mLD group (0.78% ± 0.095 of total live DRG cells for sHD compared to 0.42% ± 0.104 for the mLD, p=0.0015, p=13-15). Although not significantly greater compared to baseline, the frequency of LAP^+^ CD4^+^ T cells out of total live DRG cells at day 3 for the mLD was significantly greater than the frequency of LAP^+^ CD4^+^ T cells from the sHD group (0.59% ± 0.130 of total live DRG cells for mLD compared to 0.25% ± 0.076 for the sHD, p=0.0152, p=9) **(Fig 2D)**.

Even though the predominant CD4^+^ T cell subtypes induced by PTX in the DRG were anti-inflammatory, pro-inflammatory CD4^+^ T cells emerged after administration of the sHD and mLD PTX regimens **(Fig 2E, F)**. Following the same temporal pattern as the anti-inflammatory subtypes, the frequency of IL-17A^+^ CD4^+^ T cells for the mLD group was greatest at day 3 (>2.2-fold: 0.38% ± 0.043 of total live DRG cells at baseline to 0.84% ± 0.113 at day 3, p=0.0296, n=9), which was significantly higher than the frequency induced by the sHD (0.42% ± 0.147, p=0.0106, n=9) **(Fig 2E)**. Although not significant from baseline, the frequency of interferon (IFN)-γ ^+^ CD4^+^ T cells for the mLD group was highest at day 3 (0.45% ± 0.073), which was significantly higher than the sHD group (0.19% ± 0.049, p=0.0216, n=9) **(Fig 2F)**. In contrast, the frequency of IFN-γ^+^ CD4^+^ T cells for the sHD was highest at day 14 (>2.3-fold: 0.21% ± 0.038 of total live DRG cells at baseline to 0.50% ± 0.116 at day 14, p=0.0335, n=9-13), which was significantly higher than the mLD group (0.30% ± 0.07, p=0.0337, n=13-15) **(Fig 2F)**.

### mLD PTX induces a faster and more diverse CD8^+^ T cell response in the DRG compared to sHD PTX

Since CD4^+^ T cells are known to modulate CD8^+^ T cell responses[27], and previous studies have shown a role for CD8^+^ T cells in PTX-induced mechanical hypersensitivity[26; 57], we quantified the frequencies of CD8^+^ T cells in the DRG at baseline and after the administration of the two PTX dosing regimens. Like the CD4^+^ T cell response, the mLD PTX regimen induced CD8^+^ T cells earlier at day 3 **(Fig 3A-E)**. There were significant increases for IL-10 (>2.6-fold: 0.24% ± 0.018 of total live DRG cells at baseline to 0.64% ± 0.101 at day 3, p=0.0028, n=9) **(Fig 3A)**, LAP (3.2-fold: 0.21% ± 0.036 of total live DRG cells at baseline to 0.68% ± 0.107 at day 3, p<0.0001, n=9) **(Fig 3B)**, FoxP3 (>2.4-fold: 0.22% ± 0.046 of total live DRG cells at baseline to 0.54% ± 0.081 at day 3, p=0.0046, n=9) **(Fig 3C)**, and IL-4 (>2.6-fold: 0.16% ± 0.029 of total live DRG cells at baseline to 0.42% ± 0.041 at day 3, p=0.0016, n=9) **(Fig 3D)** CD8^+^ T cells. Again, these frequencies at day 3 were significantly higher for the mLD compared to the sHD for IL-10 (0.24% ± 0.018, p=0.0001, n=9) **(Fig 3A)**, LAP (0.16% ± 0.028, p<0.0001, n=9) **(Fig 3B)**, FoxP3 (0.15% ± 0.029, p<0.0001, n=9) **(Fig 3C)**, and IL-4 (0.12% ± 0.026, p<0.0001, n=9) **(Fig 3D)**. In addition to these anti-inflammatory CD8^+^ T cell populations, the mLD regimen induced a pro-inflammatory IL-17A^+^ CD8^+^ T cell response at day 3 (>2.1-fold: 0.3% ± 0.041 of total live DRG cells at baseline to 0.65% ± 0.103 at day 3, p=0.0010, n=9), which was significantly greater than the sHD regimen (0.22% ± 0.031 at day 3, p<0.0001, n=9) **(Fig 3E)**.

**Figure 3.**
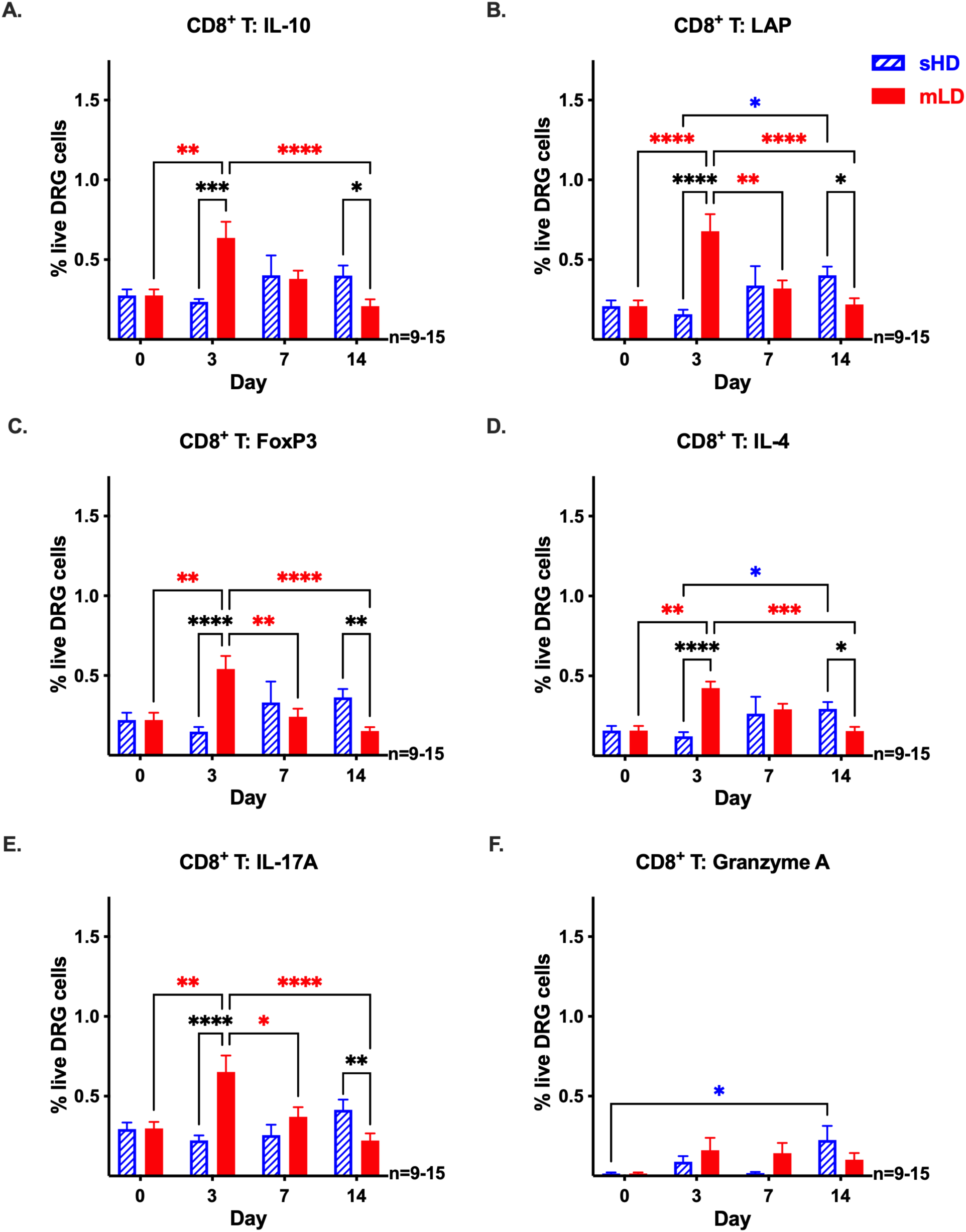
The mLD PTX regimen induces a faster and more diverse CD8^+^ T cell response in the DRG than the sHD regimen. Frequencies of **(A)** IL-10, **(B)** LAP, **(C)** FoxP3, **(D)** IL-4, **(E)** IL-17A, and **(F)** granzyme A CD8^+^ T cells out of total live DRG cells on day 0 and after sHD (blue) or mLD (red) PTX regimens. Significance was determined by 2-way ANOVA with Tukey’s multiple comparisons test (*p<0.05, **p<0.01, ***p<0.001, ****p<0.0001, n=9-15/regimen). Blue (sHD) and red (mLD) significance asterisks compare CD8^+^ T cell frequencies within each dosing regimen. Black significance asterisks compare CD8^+^ T cell frequencies between sHD and mLD PTX regimens.

This robust CD8^+^ T cell response induced by the mLD PTX regimen was further supported by IL-2 production from CD4^+^ T cells **(S4A Fig)** and CD8^+^ T cells **(S4B Fig)**. In contrast, the sHD regimen did not induce much of an IL-2 T cell response **(S4A, B Fig)** or anti-inflammatory CD8^+^ T cells in the DRG **(Fig 3A-D)**. The CD8^+^ T cell response for the sHD was primarily pro-inflammatory subtypes **(Fig 3E, F)**. There was a significant increase in granzyme A^+^ CD8^+^ T cells at day 14 (>15-fold: 0.015% ± 0.007 of total live DRG cells at baseline to 0.23% ± 0.089 at day 14, p=0.0463, n=9-13) **(Fig 3E)**. In addition, the sHD induced significantly more IL-17A^+^ CD8^+^ T cells at day 14 (0.42% ± 0.064, p=0.0095, n=13) compared to the mLD (0.22% ± 0.045, n=15) **(Fig 3F)**. This predominant pro-inflammatory CD8^+^ T cells response in the sHD regimen might in part explain why the sHD mice were more hypersensitive at day 14 than the mLD group.

### CD3^−^ cells promote T cell function after administration of PTX

To further support this strong PTX-induced T cell response, we evaluated a potent T cell growth factor, IL-2. We detected IL-2 production by CD3^−^ cells in the DRG on day 3 for both sHD and mLD treated mice **(Fig 4A)**. There was a significant increase in CD3^−^ IL-2 production on day 7 for the mLD group (>15-fold: 2.45% ± 0.270 of total live DRG cells at baseline to 38.79% ± 14.34 at day 7, p=0.0106, n=9), which was significantly greater than the frequency induced by the sHD regimen (10.55% ± 5.53 at day 7, p=0.0151, n=9) **(Fig 4A)**. In addition to IL-2, we detected granzyme A production in CD3^−^ cells after sHD and mLD PTX treatment **(Fig 4B)**. The frequency of granzyme A^+^ CD3^−^ cells was significantly greater at day 7 in the mLD compared to the sHD (10-fold: 36.05% ± 14.98 of total live DRG cells for mLD compared to 3.57% ± 2.17 for the sHD, p=0.0222, n=9) **(Fig 4B)**. These CD3^−^ cells producing IL-2 and granzyme A are likely natural killer (NK) cells, and together, this suggest that PTX may promote NK cell function to support T cell function.

**Figure 4.**
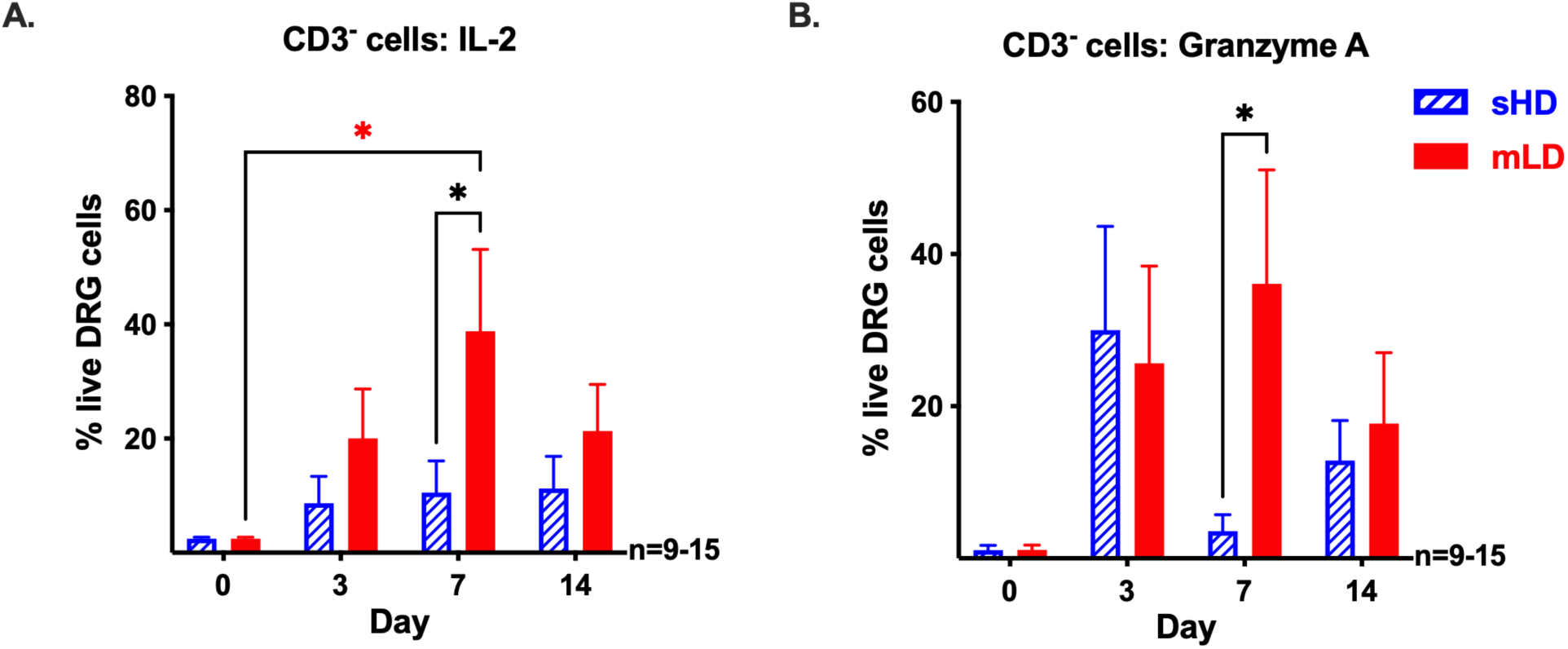
CD3^−^ cells produce IL-2 and granzyme A after PTX. Frequencies of **(A)** IL-2, **(B)** granzyme A CD3^−^cells out of total live DRG cells on day 0 and after sHD (blue) or mLD (red) PTX regimens. Significance was determined by 2-way ANOVA with Tukey’s multiple comparisons test (*p<0.05, n=9-15/regimen). Red significance asterisk compares CD3^−^ cell frequency after mLD PTX to day 0. Black significance asterisks compare CD3^−^ cell frequencies between sHD and mLD PTX regimens.

**Figure 5.**
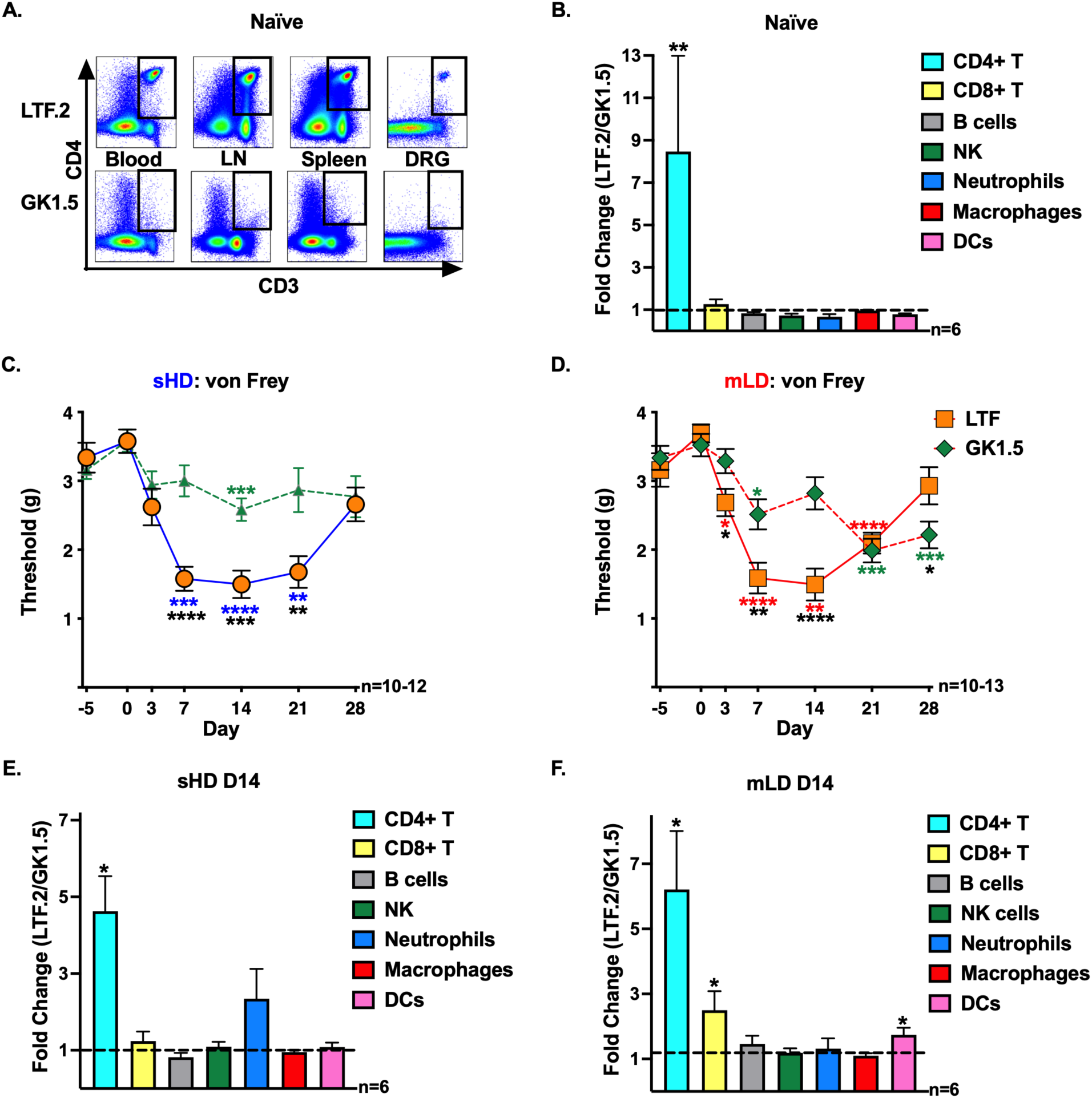
CD4^+^ T cell depletion reduces PTX-induced mechanical hypersensitivity. **(A)** Flow cytometric pseudocolor dot plots showing CD3^+^ CD4^+^ T cells in the blood, lymph nodes (LN), spleen, and DRG from naïve female mice injected with LTF.2 (isotype control antibody, top) or GK1.5 (CD4^+^ T cell depleting antibody, bottom). **(B)** Fold changes of CD45^+^ immune cell populations from DRG represent the ratio of immune cell frequencies in naïve LTF.2 control mice to those in GK1.5-treated (CD4^+^ T cell depleted) mice. The dashed line at 1 represents no change. Values above the dashed line indicate an increase in the LTF.2 control mice or a decrease in the GK1.5 depleted mice. Asterisks denote statistically significant differences in cell frequencies between naïve LTF.2 and GK1.5-treated mice within each cell subset, calculated using an unpaired t-test with Welch’s correction (**p<0.01). **(C, D)** Mechanical hypersensitivity (tactile threshold in grams) was assessed in CD4^+^ T cell sufficient (LTF.2) or CD4^+^ T cell deficient (GK1.5) mice with von Frey filaments prior to antibody injection day (D) -5, prior to PTX (D 0) to determine baseline (BL) values, and after **(C)** sHD or **(D)** mLD PTX on D 3, 7, 14, 21, and 28. Significance was determined by 2-way repeated measures ANOVA with Tukey’s multiple comparisons test (*p<0.05, **p<0.01, ***p<0.001, ****p<0.0001, n=9-13/regimen). Blue (sHD) and red (mLD) significance asterisks compare hypersensitivity after PTX to BL within each LTF.2 group. Green significance asterisks compare hypersensitivity after PTX to BL within each GK1.5 group. Black significance asterisks compare hypersensitivity values between LTF. 2 and GK1.5 groups. **(E, F)** Fold changes of CD45^+^ immune cell populations from DRG represent the ratio of immune cell frequencies from D 14 **(E)** sHD or **(F)** mLD LTF.2 control mice to those in GK1.5-treated (CD4^+^ T cell depleted) mice. Asterisks denote statistically significant differences in cell frequencies within each cell subset, calculated using an unpaired t-test with Welch’s correction (*p<0.05).

**Figure 6.**
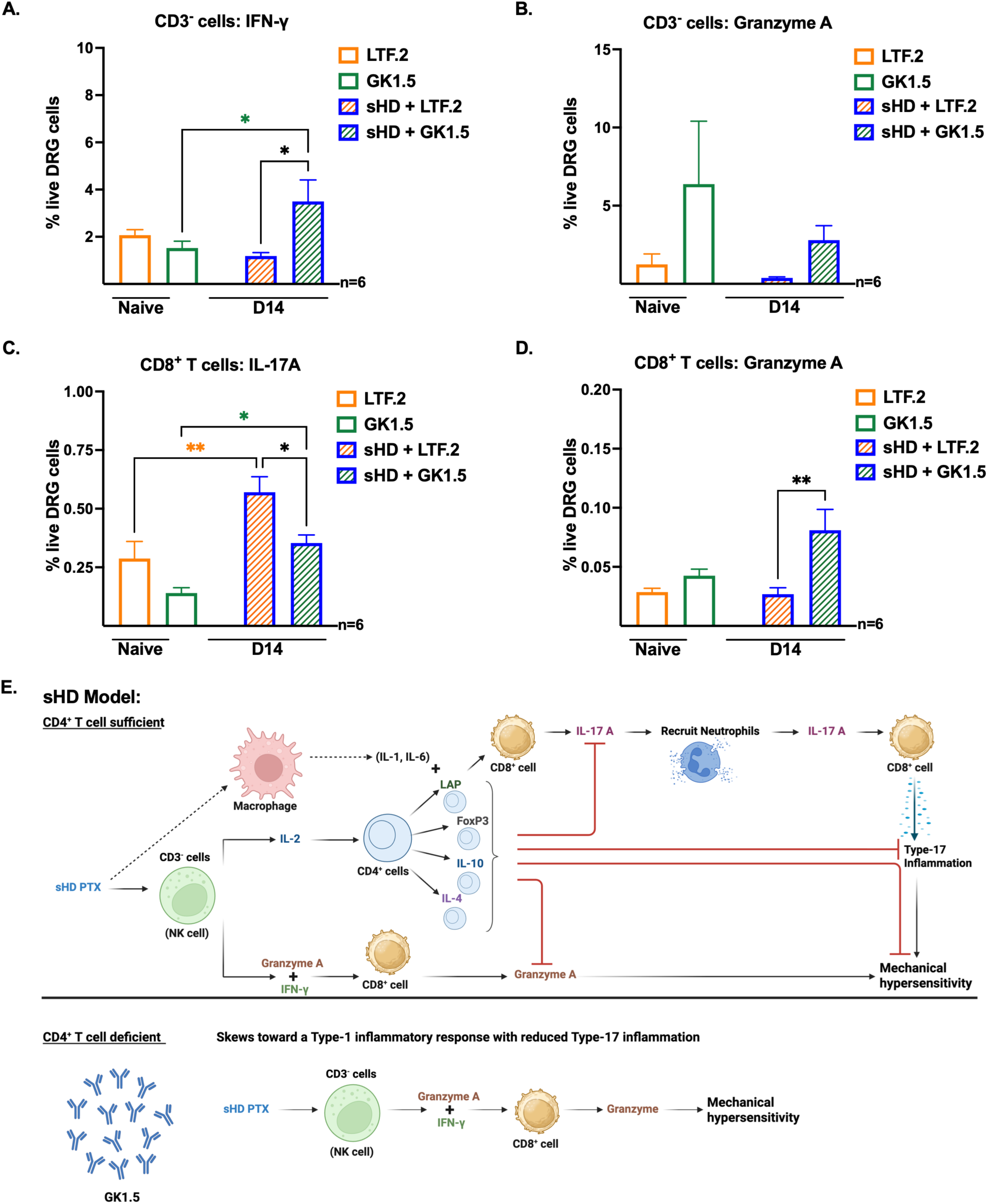
sHD PTX skews toward a Type-1 inflammatory response with reduced Type-17 inflammation in CD4^+^ T depleted mice. Frequencies of **(A)** CD3^−^ IFN-γ^+^ cells, **(B)** CD3^−^ granzyme A^+^ cells, **(C)** CD8^+^ IL-17A^+^ cells, **(D)** CD8^+^ granzyme A^+^ cells out of total live cells in DRG from naïve mice treated with LTF.2 (white bar with orange border) or GK1.5 (white bar with green border), and from sHD day (D) 14 mice treated with LTF.2 (bar with orange diagonal lines and blue border) or GK1.5 (bar with green diagonal lines and blue border). Significance was determined by 1-way ANOVA with Tukey’s multiple comparisons test (*p<0.05, **p<0.01, n=6). Orange (LTF.2) and green (GK1.5) significance asterisks compare cell frequencies between naïve and sHD D 14 within each antibody group. Black significance asterisks compare cell frequencies between LTF.2 and GK1.5 groups. **(E)** Model for immune cell function and PTX-induced hypersensitivity after sHD PTX regimen in CD4^+^ T sufficient (top) or CD4^+^ T cell deficient (bottom) mice. Dashed arrows indicate information obtain from literature. Words in parentheses indicate the inferred cell type or cytokines based on cell characteristics. Black arrowheads indicate secretion or promotion of activity, whereas red flatheads (⊥) indicate inhibition or reduced activity. Graphic created by Biorender.

### CD4^+^ T cell depletion modulates immune cell frequencies in the DRG and reduces PTX-induced mechanical hypersensitivity

To further understand the role of CD4^+^ T cells in PTX-induced peripheral neuropathy, we depleted CD4^+^ T cells in adult female C57BL6/J mice by injecting an antibody against the CD4 receptor (GK1.5). CD4^+^ T cell depletion was verified by loss of CD3^+^ CD4^+^ T cells in blood, lymph nodes, spleen, and DRG after GK1.5 injection, and the maintenance of CD3^+^ CD4^+^ T cells after LTF.2 antibody control injection **(Fig 5A)**. The LTF.2 control mice had 8.5-fold more CD4^+^ T cells (p=0.0034, n=6) than the CD4^+^ T cell depleted mice **(Fig 5B).** All other immune populations (CD8^+^ T cells, B cells, NK cells, neutrophils, macrophages, and DCs) were relatively unchanged after CD4^+^ T cell depletion **(Fig 5B)**. We assessed mechanical hypersensitivity on day -5 to obtain von Frey values prior to GK1.5 or LTF.2 injection (day -4) **(Fig 5C, D)**. After depleting CD4^+^ T cells, the mice were tested again on day 0 to determine the extent CD4^+^ T cells contribute to mechanical sensitivity in the absence of chemotherapy. CD4^+^ T cell depletion or injection of the LTF.2 control antibody did not change mechanical sensitivity at day 0 compared to day -5 **(Fig 5C, D)**. However, depleting CD4^+^ T cells drastically reduced PTX-induced mechanical hypersensitivity for both the sHD (day 7: 3.000 ± 0.226 for GK1.5 and 1.579 ± 0.174 for LTF.2, p<0.0001; day 14: 2.583 ± 0.162 for GK1.5 and 1.497 ± 0.199 for LTF.2, p=0.0005; day 21: 2.867 ± 3.19 for GK1.5 and 1.677 ± 0.231 for LTF.2, p=0.0070; n=10-12) **(Fig 5C)** and mLD (day 3: 3.288 ± 0.178 for GK1.5 and 2.685 ± 0.200 for LTF.2, p=0.0357; day 7: 2.513 ± 0.220 for GK1.5 and 1.586 ± 0.224 for LTF.2, p=0.0078; day 14: 2.819 ± 0.235 for GK1.5 and 1.493 ± 0.232 for LTF.2, p=0.0006; n=10-13) **(Fig 5D)** regimens with a significantly greater reduction in hypersensitivity with the sHD regimen compared to the mLD regimen (day 21: 1.987 ± 0.172 for mLD + GK1.5 and 2.867 ± 0.319 for sHD + GK1.5, p=0.0267; n=12-13) **(S5 Fig)**. The sHD + GK1.5 group was only significantly more hypersensitive than baseline on day 14 (2.583 ± 0.162, p=0.0003, n=12) **(Fig 5C)** while the mLD + GK1.5 group was significantly more hypersensitive than baseline at day 7 (2.513 ± 0.220, p=0.0127), D 21 (1.987 ± 0.172, p=0.0004) and day 28 (2.215 ± 0.195, p=0.0006) **(Fig 5D)**.

Although PTX caused hypersensitivity in the LTF.2 controls, the hypersensitivity values at day 3 for the sHD group without LTF.2 was less severe than the sHD (1.656 ± 0.131 for sHD and 2.62 ± 0.267 for sHD + LTF.2, p=0.0279, n=10-20), possibly suggesting an Fc antibody effect **(S6A Fig)**. When we evaluated the immune cell populations in the DRG, we found that the naïve plus LTF.2 group had a 7.25-fold decrease in the frequency of neutrophils on day 0 (prior to PTX injection) (0.087% ± 0.024 neutrophils out of total live DRG cells in naïve compared to 0.012% ± 0.002 in naïve + LTF.2 control, p=0.0053, n=6) **(S6B Fig)**, suggesting a decrease or delay in the inflammatory response as neutrophils are usually the first responders to injury. When we evaluated the frequency of neutrophils on day 14, we found that there was no difference with or without LTF.2 **(S6B Fig)**. However, compared to baseline levels, there was a significant decrease in the frequency of neutrophils only after mLD PTX (0.087% ± 0.024 neutrophils out of total live DRG cells in naïve compared to 0.012% ± 0.003 in mLD at D 14, p=0.0052, n=6). In contrast, neutrophils were maintained after treatment with sHD PTX (0.087% ± 0.024 neutrophils out of total live DRG cells in naïve compared to 0.053% ± 0.012 in sHD at day 14, p=0.3751, n=6-10), suggesting neutrophils may contribute to mechanical hypersensitivity with the sHD regimen.

Since CD4^+^ T cells impact the survival and function of other immune cells[71], we evaluated the fold change of CD45^+^ immune cells in the DRG after CD4^+^ T cell depletion from day 14 sHD **(Fig 5E)** or mLD **(Fig 5F)** PTX-treated mice. After sHD treatment, the CD4^+^ T cell depleted mice had significantly fewer CD4^+^ T cells (4.6-fold, p=0.0115, n=6) compared to CD4^+^ T cell sufficient mice **(Fig 5E)**. In contrast, the CD4^+^ T cell depleted mice that received the mLD regimen had fewer CD4^+^ T cells (6.2-fold, p=0.0418, n=6), CD8^+^ T cells (2.5-fold, p=0.0410, n=6), and DCs (1.7-fold, p=0.0434, n=6) compared to CD4^+^ T cell sufficient mice **(Fig 5F)**. This data suggests that CD4^+^ T cell depletion modulates the immune microenvironment in the DRG, which together could impact mechanical hypersensitivity.

### sHD PTX skews immune cells toward a Type-1 inflammatory response in CD4^+^ T depleted mice

To determine how CD4^+^ T cell depletion impacts PTX-induced mechanical hypersensitivity, we evaluated cytokine production from CD3^−^ cells **(Fig 6A, B)** and CD8^+^ T cells **(Fig 6C, D)**. We found that after CD4^+^ T cell depletion, sHD PTX increased IFN-γ producing CD3^−^ cells at day 14 (>2.3-fold: 1.53% ± 0.285 of total live DRG cells at baseline to 3.5% ± 0.913 at day 14, p=0.0497, n=6) **(Fig 6A)**. Moreover, the frequency of CD3^−^ IFN-γ^+^ cells at day 14 was significantly greater in the CD4^+^ T cell depleted mice compared to the LTF. 2 controls (1.19% ± 0.144, p=0.0178, n=6) **(Fig 6A)**. Although there was no significant increase in granzyme A production in CD3^−^ cells **(Fig 6B)**, the maintenance of granzyme A production in CD3^−^ cells along with an increase in IFN-γ supports a skew toward Type-1 innate immunity after CD4^+^ T cell depletion. In addition, CD4^+^ T cell depletion prevented the sHD PTX-induced increase in CD8^+^ T cell IL-17A production. There was a significant increase in IL-17A^+^ CD8^+^ T cells at day 14 in the LTF.2 control mice (almost 2-fold: 0.29% ± 0.073 of total live DRG cells at baseline to 0.57% ± 0.066 at day 14, p=0.0066, n=6), which was significantly greater than the CD4^+^ T cell depleted mice (0.35% ± 0.035, p=0.0441, n=6) **(Fig 6C)**. Since the innate Type-1 response was intact, we did see an increase in granzyme A production in CD8^+^ T cells in the CD4^+^ T cell depleted mice (2-fold: 0.04% ± 0.006 of total live DRG cells at baseline to 0.08% ± 0.018 at day 14, p=0.0514, n=6), which was significantly greater than the LTF. 2 controls (0.03% ± 0.005, p=0.0043, n=6) **(Fig 6D)**. This increase in granzyme A^+^ CD8^+^ T cells in the CD4^+^ T cell depleted mice might contribute to the slight hypersensitivity detected at day 14.

Based on published literature and our data from the CD4^+^ T cell sufficient and deficient mice, we developed a model of immune cell function and PTX-induced mechanical hypersensitivity after the sHD regimen **(Fig 6E)**. In CD4^+^ T cell sufficient mice, a sHD of PTX activates CD3^−^ cells (most likely NK cells) to secrete IFN-γ, granzyme A, and IL-2. IFN-γ and granzyme A production induces CD8^+^ T cells to secrete granzyme A, which we propose causes mechanical hypersensitivity as granzymes has been shown to damage neurons[42]. The IL-2 produced by the CD3^−^ cell (most likely NK cells) will then promote the survival, activation, and differentiation of CD4^+^ T cells to LAP, FoxP3, IL-10, and IL-4 subtypes. As mentioned previously, LAP can either promote pro- or anti-inflammatory immune responses, depending on other cytokines in the local milieu. It is well established that PTX induces IL-1 and IL-6 production in the DRG[4] with the macrophage as the most likely producer[69]. IL-1 and IL-6 combined with LAP from the CD4^+^ T cell can support the activation and differentiation of IL-17A producing CD8^+^ T cells[32]. IL-17A production by CD8^+^ T cells can recruit neutrophils[48] to the DRG, which can produce more IL-17A, providing a feed-forward loop (to further enhance IL-17A production from CD8^+^ T cells). This Type 17 inflammation will promote mechanical hypersensitivity[34]. Now, it is known that an injury normally produces both pro- and anti-inflammatory immune responses to promote repair and prevent excessive tissue damage. Here, the anti-inflammatory FoxP3, IL-10, and IL-4 producing CD4^+^ T cells will suppress Type-17 inflammation and granzyme A production, leading to the resolution of PTX-induced mechanical hypersensitivity. In the absence of CD4^+^ T cells, the immune response skews toward a Type-1 inflammatory response instead of a Type-17 inflammatory response. More specifically, the sHD of PTX only activates the CD3^−^ cells (most likely NK cells) to secrete IFN-γ and granzyme A. IFN-γ and granzyme A production will support granzyme A^+^ CD8^+^ T cells[43], which will induce some mechanical hypersensitivity as we saw in the CD4^+^ T cell depleted mice. Overall, the level of PTX-induced mechanical hypersensitivity is much lower in the CD4^+^ T cell depleted mice as Type-17 inflammation is significantly reduced in the absence of CD4^+^ T cells.

### mLD PTX regimen promotes LAP suppressive state in CD4^+^ T depleted mice

To determine how CD4^+^ T cell depletion impacts PTX-induced mechanical hypersensitivity after the mLD treatment, we evaluated cytokine production from CD3^−^ cells **(Fig 7A-C)** and CD8^+^ T cells **(Fig 7D, S7 Fig)**. At day 14, we found that CD3^−^ IL-17A^+^ cells (5.65% ± 0.716 for GK1.5 and 1.57% ± 0.240 for LTF.2, p<0.0001; n=6) **(Fig 7A)** and CD3^−^ LAP^+^ cells (9.32% ± 1.059 for GK1.5 and 2.85% ± 0.466 for LTF.2, p=0.0008; n=6) **(Fig 7B)** were significantly greater in the CD4^+^ T cell depleted mice than the LTF.2 control mice. Moreover, the frequency of CD3^−^ LAP^+^ cells were significantly decreased in the LTF.2 control mice (>3.4-fold: 9.90% ± 0.919 of total live DRG cells at baseline to 2.85% ± 0.466 at day 14, p=0.0003, n=6), but remained constant in CD4^+^ T cell depleted mice (9.67% ± 1.299 of total live DRG cells at baseline to 9.32% ± 1.059 at day 14) **(Fig 7B)**. In addition, CD4^+^ T cell depletion prevented an increase in granzyme A production in both CD3^−^ cells **(Fig 7C)** and CD8^+^ T cells **(Fig 7D)**. There was a significant increase in the frequency of CD3^−^ granzyme A^+^ cells in the LTF. 2 control mice (>30-fold: 1.24% ± 0.670 of total live DRG cells at baseline to 38.12% ± 13.27 at day 14, p=0.0063, n=6), which was significantly more than the CD4^+^ T cell depleted mice (0.52% ± 0.084, p=0.0053, n=6) **(Fig 7C)**. Similarly, there was a significant increase in the frequency of CD8^+^ granzyme A^+^ cells in the LTF. 2 control mice (>12.6-fold: 0.03% ± 0.003 of total live DRG cells at baseline to 0.38% ± 0.106 at day 14, p=0.0007, n=6), which was significantly more than the CD4^+^ T cell depleted mice (0.03% ± 0.003, p=0.0008, n=6) **(Fig 7D)**. In addition, CD4^+^ T cell depleted mice had significantly fewer FoxP3^+^, LAP^+^, and IL-4^+^ CD8^+^ T cells than the LTF.2 controls **(S7A, B Fig).**

**Figure 7.**
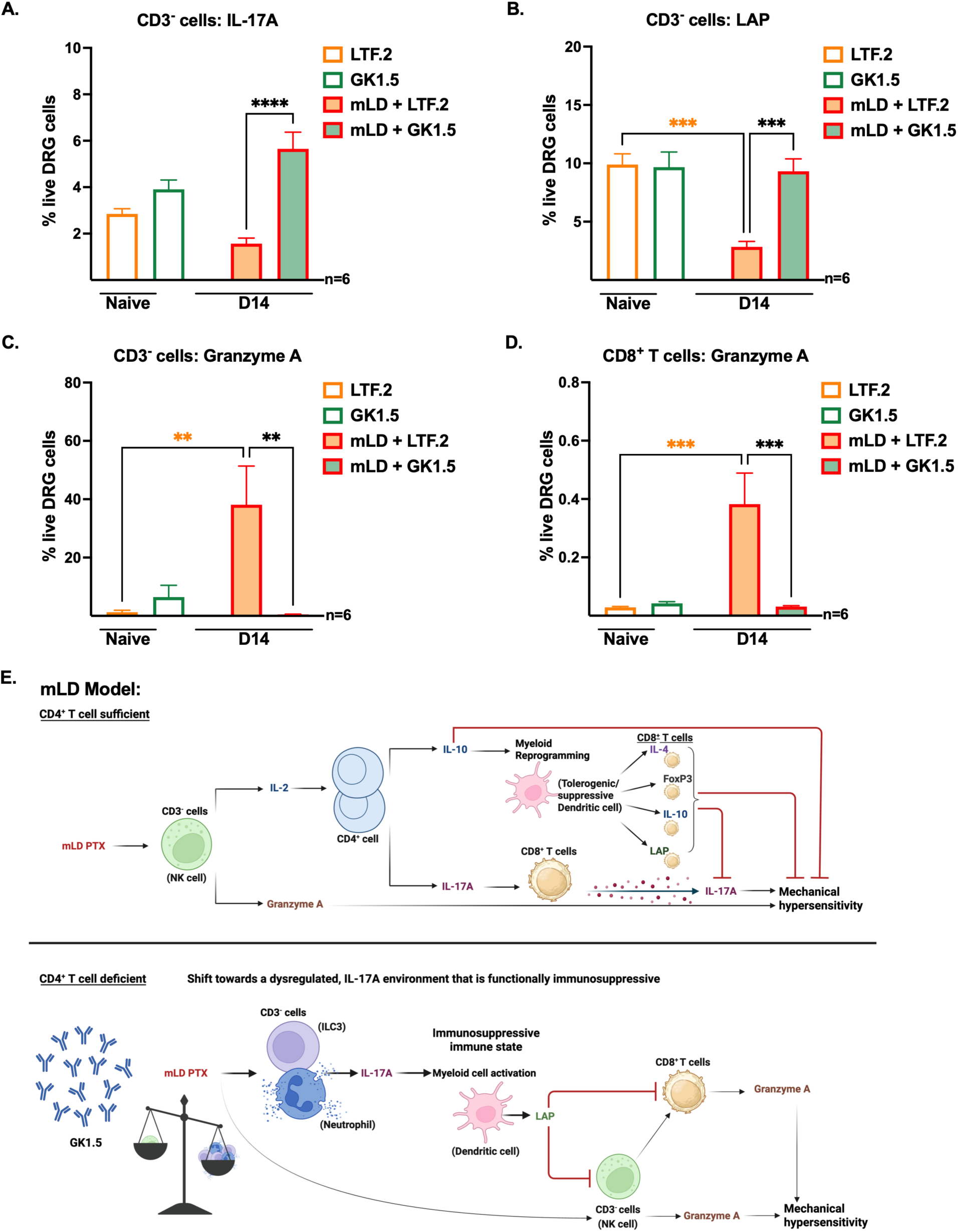
mLD PTX shifts the response toward an LAP suppressive state in CD4^+^ T depleted mice. Frequencies of **(A)** CD3^−^ IL-17A^+^ cells, **(B)** CD3^−^ LAP^+^ cells, **(C)** CD3^−^ granzyme A^+^ cells, **(D)** CD8^+^ granzyme A^+^ cells out of total live cells in DRG from naïve mice treated with LTF.2 (white bar with orange border) or GK1.5 (white bar with green border), and from mLD day (D) 14 mice treated with LTF.2 (orange bar with red border) or GK1.5 (green bar with red border). Significance was determined by 1-way ANOVA with Tukey’s multiple comparisons test (**p<0.01, ***p<0.001, ****p<0.0001, n=6). Orange significance asterisks compare cell frequencies between naïve and mLD D 14 within LTF.2 antibody group. Black significance asterisks compare cell frequencies between LTF.2 and GK1.5 groups. **(E)** Model for immune cell function and PTX-induced hypersensitivity after mLD PTX regimen in CD4^+^ T sufficient (top) or CD4^+^ T cell deficient (bottom) mice. Words in parentheses indicate inferred cell type based on cell characteristics. Black arrowheads indicate secretion or promotion of activity, whereas red flatheads (⊥) indicate inhibition or reduced activity. Graphic created by Biorender.

Like the sHD regimen, we propose that the mLD regimen induces mechanical hypersensitivity via granzyme A and IL-17A; however, the cellular pathways are different **(Fig 7E)**. Like the sHD, the mLD PTX activates CD3^−^ cells (most likely NK cells) to produce granzyme A and IL-2. CD3^−^ granzyme A production can cause PTX-induced mechanical hypersensitivity directly while IL-2 production promotes the CD4^+^ T cell response. The first main CD4^+^ T cell population induced by mLD PTX is the IL-17A^+^ producing CD4^+^ T cells, which can activate IL-17A^+^ CD8^+^ T cells to induce mechanical hypersensitivity. The other major CD4^+^ T cell population secretes anti-inflammatory IL-10, acting on DCs to induce differentiation to a tolerogenic/suppressive phenotype. Now, when suppressive DCs engage with CD8^+^ T cells, the DC will promote anti-inflammatory CD8^+^ T cell subtypes (IL-4, FoxP3, IL-10, and LAP). Together, these anti-inflammatory CD4^+^ and CD8^+^ T cells will suppress PTX-induced mechanical hypersensitivity. In the absence of CD4^+^ T cells, mLD PTX will induce a smaller CD3^−^ granzyme A^+^ (most likely NK cells) population, and a much larger population of IL-17A producing CD3^−^ cells (most likely innate lymphoid cells 3 (ILC3) and/or neutrophils). During this initial immune response (day 0 through day 14), the smaller CD3^−^ granzyme A^+^ cells will induce some PTX-induced mechanical hypersensitivity while the larger IL-17A^+^ CD3^−^ cell population will promote an immunosuppressive state. Here, IL-17A production activates a CD3^−^ cell (most likely a DC) to secrete LAP[53]. LAP will inhibit granzyme A production from both CD3^−^ cells (NK cells) and CD8^+^ T cells[61], reducing PTX-induced mechanical hypersensitivity.

Collectively, the dose and frequency of PTX induces distinct immune cell populations, which contribute to the severity of mechanical hypersensitivity and the induction of the ATF-3 driven transcriptional neuronal stress pathway. PTX induces both pro- and anti-inflammatory populations with the pro-inflammatory ones promoting CIPN and anti-inflammatory ones controlling/suppressing CIPN. In the presence of CD4^+^ T cells, CIPN likely develops due to Type-17 inflammation, but in the absence of CD4^+^ T cells, the immune environment shifts toward a Type-1 inflammatory response for sHD, and a dysregulated Type-17, immunosuppressive LAP response for mLD. Overall, the shifts in immune cell function in the absence of CD4^+^ T cells is less inflammatory resulting in significantly decreased mechanical hypersensitivity.

## DISCUSSION

The present study demonstrates that PTX dose does not directly correlate with the magnitude or duration of PTX-induced mechanical hypersensitivity or the activation of DRG stress markers like ATF-3, challenging the traditional view of CIPN as a purely cumulative, dose-dependent toxicity. Instead, our data indicate that PTX dosing intensity and frequency shape the immune landscape within the DRG, particularly the CD4^+^ T cell compartment, which in turn orchestrates innate and adaptive immune responses. These findings support a model in which CIPN reflects a neuroimmune toxicity driven by context-dependent immune programming rather than linear drug accumulation.

Importantly, PTX does not introduce a novel exogenous antigen. Rather, chemotherapy-induced tissue injury most likely leads to the release of endogenous self-antigens derived from stressed or damaged cells. These antigens, often modified by oxidative or metabolic stress, are accompanied by danger-associated molecular patterns (DAMPs), including HMGB1, ATP, and uric acid. DAMPs are recognized primarily by macrophages and DCs, leading to cytokine production that critically shapes CD4^+^ T cell differentiation. While antigen dose influences T cell receptor (TCR) signal strength and can bias helper polarization[2; 16; 37], accumulating evidence indicates that the DAMP-conditioned cytokine milieu is the dominant determinant of whether CD4^+^ T cells adopt a broad polyfunctional repertoire or a more restricted helper phenotype[46]. Thus, the “danger context,” rather than antigen quantity alone, governs adaptive CD4^+^ T cell diversification following PTX exposure.

Our comparative PTX dosing analysis of a single, high dose (sHD) versus multiple low doses (mLD, higher cumulative dose) illustrates this principle. The mLD PTX regimen induces an early T cell response by day 3, characterized by both pro- and anti-inflammatory T cell subsets, whereas the sHD PTX regimen elicits a delayed response emerging at day 14. Moreover, the sHD regimen induces greater mechanical hypersensitivity and the activation of the ATF3-mediated neuronal stress pathway, suggesting more extensive tissue injury and DAMP release[10; 11; 31]. Consistent with the danger hypothesis, the sHD PTX regimen promotes broader CD4^+^ T cell diversification, likely reflecting exposure to a more heterogeneous DAMP environment. Mechanistically, HMGB1 enhances Th1 polarization[49], extracellular ATP promotes Th17 differentiation[21], and uric acid drives Th2 responses[25]. Greater tissue stress would therefore be expected to expand multiple helper subsets simultaneously, generating T cell diversity. In contrast, PTX-induced mechanical hypersensitivity is less severe in the mLD group with fewer ATF3^+^ DRG neurons, likely generating fewer DAMPs, which produces a more restricted CD4^+^ T cell profile (IL-17A and IL-10). Future studies will quantify intraepidermal nerve fibers in the hindpaw and characterize DAMPs after PTX exposure as interventions targeting DAMPs may be effective at shifting toward a neuroprotective immune response. In addition to DAMPs, repeated antigenic stimulation also influences CD4^+^ T cell fitness. Both in vitro and in vivo studies demonstrate that repeated stimulation of effector CD4^+^ T cells reduces proliferative capacity and cytokine production[18]. Thus, dosing frequency in the mLD PTX regimen may modulate not only the magnitude but also the durability of helper responses as the mLD CD4^+^ T cell response decreases with additional PTX injections.

In contrast to CD4^+^ T cells, CD8^+^ T cells are more dependent on sustained antigen exposure and TCR signal strength for optimal differentiation[3; 46]. This distinction likely explains the stronger, clonally diverse CD8^+^ T cell expansion observed in the mLD regimen, which may provide sufficient antigen persistence to drive differentiation. Collectively, these findings indicate that mLD PTX induces selective immune shaping—favoring CD8^+^ T cell diversification with limited CD4^+^ breadth—whereas sHD PTX induces diversity characterized by expanded CD4 helper heterogeneity but weaker, more polarized CD8^+^ responses.

Our data further implicate IL-17A signaling as a key mediator of PTX-induced hypersensitivity. IL-17 receptor expression is widely detected in DRG neurons and cultured sensory neurons[47; 51]. IL-17A contributes to neuropathic pain across multiple models—including chronic constriction injury, partial sciatic ligation, neuritis, and CIPN[34; 40]—and directly enhances neuronal excitability in vitro[47]. IL-17A-deficient mice exhibit reduced mechanical hypersensitivity and decreased T cell and macrophage infiltration in injured nerves[23; 24; 51]. In our study, the sHD regimen selectively expanded the IL-17A^+^ CD8^+^ T cell subset, while the mLD regimen fostered a more heterogeneous environment characterized by the involvement of both IL-17A producing CD4^+^ and CD8^+^ T cells. A previous study suggested that IL-17-driven CIPN occurs independently of T cells; however, T cells were assessed only at 7 days post-sHD PTX[34]. Thus, early IL-17A responses after PTX may be glial- or neutrophil-derived, whereas later IL-17A likely reflects adaptive T cell contributions, consistent with the delayed CD4^+^ expansion observed after the sHD PTX regimen. These findings support a model in which IL-17A acts both directly on neurons and indirectly by recruiting and activating immune cells within the DRG microenvironment.

Granzyme A may represent an additional mechanism linking cytotoxic immune cell activation to neuronal injury. Unlike granzyme B, granzyme A induces caspase-independent programmed cell death through mitochondrial disruption and reactive oxygen species generation[38]. Cytotoxic T lymphocyte-derived granzyme A can trigger neurite retraction via thrombin receptor activation[60] and promote inflammatory cytokine production, specifically by cleaving pro-IL-1β [17; 39; 58]. These findings suggest concentration-dependent effects: low granzyme A levels amplify inflammation, whereas higher levels promote neuronal damage. Thus, granzyme A may contribute to both inflammatory amplification in both the sHD and mLD PTX regimens but structural neuronal injury in only the sHD PTX regimen.

Depletion of CD4^+^ T cells significantly reduced PTX-induced mechanical hypersensitivity, with a stronger effect in the sHD PTX group. In addition to CD4^+^ T cells, our data suggests that neutrophils may contribute to PTX-induced hypersensitivity, particularly in the sHD PTX group. Neutrophils contribute to CIPN pathogenesis[30; 64], and their depletion suppresses IL-17A-driven inflammation[70]. Similar CD4^+^ T cell–dependent neutrophil amplification has been observed in lung ischemia-reperfusion injury and pneumococcal pneumonia, where CD4^+^ depletion reduced neutrophil infiltration and IL-17A levels[54; 68]. These parallels suggest that CD4^+^ T cells may amplify myeloid-driven inflammation in the sHD PTX group, promoting IL-17–dominated pathology.

Conversely, in the mLD group, CD4^+^ T cell depletion primarily affected CD8^+^ T cells and DCs. CD8^+^ T cells have been shown to promote resolution of PTX-induced hypersensitivity through IL-10–dependent mechanisms[26]. Although IL-10 production was primarily shown to be macrophage derived[57], IL-10–deficient CD8^+^ T cells exhibited delayed recovery, suggesting a partial contribution. In addition to IL-10, we found other inhibitory CD8^+^ T cell populations, (IL-4, TGF-β, and FoxP3), which may contribute to resolution. Moreover, our mLD PTX regimen consisted of four injections of 2 mg/kg, whereas Krukowski et al. administered only two injections of 2 mg/kg. The increased frequency of dosing in our model likely provides more sustained antigen exposure, which is known to promote more robust CD8^+^ T cell activation and differentiation. While very few studies evaluate DCs, one suggests that an increase in immature (tolerogenic) DCs is associated with cumulative chemotherapy exposure[9] and may further contribute to a regulatory environment. Thus, in the mLD PTX group, CD8^+^ T cells may play dual roles: promoting injury via IL-17A and facilitating resolution via tolerogenic DCs driving anti-inflammatory (IL-10, IL-4, TGF-β, and FoxP3) immunoregulation.

Interestingly, CD4^+^ T cell depletion in the sHD mice decreases Type-17 inflammation but preserves the Type 1 profile in the DRG characterized by IFN-γ and granzyme A. Analogous findings in tuberculosis models demonstrate that CD4^+^ depletion maintains IFN-γ production and promotes a compensatory increase in inflammatory CD8^+^ T cells[50]. Although CD4^+^ T cell help is often required for long-term CD8^+^ T cell immunity[41], CD8^+^ cells can develop and initiate primary responses in acute inflammatory settings. In the mLD group, CD4^+^ T cell depletion results in a dysregulated IL-17 environment and promotes a LAP-associated suppressive state. Because IL-17 signaling can enhance Treg suppressive function[8], promote recruitment of MDSCs[66], and drive CD8^+^ T cell exhaustion[22], immune cell interaction that shift the environment from pro- to anti-inflammatory may critically determine the overall immune contribution to CIPN.

These findings carry important implications for patients undergoing cancer treatment with neurotoxic chemotherapies. If neuropathy is triggered early and amplified by neuroimmune cascades, dose reductions made later in a patient’s prescribed course of chemotherapy may not meaningfully reverse progression. CIPN may behave less like a linear cumulative toxicity and more like a threshold-triggered switch. Once neuroinflammatory circuits are established—particularly those involving CD4^+^ T cells, IL-17A, and granzyme A—progression may continue independent of ongoing drug exposure. Thus, reducing chemotherapy dose after symptom onset may compromise tumor control without substantially improving neuropathy risk.

These data argue for a paradigm shift from reactive dose reduction to proactive prevention. Early identification of neuronal stress biomarkers and immune signatures may enable risk stratification before irreversible neuroinflammation is established. Personalized approaches based on baseline immune profile or genetic susceptibility may prove more valuable than uniform dose modification. Mechanism-targeted interventions that skew CD4^+^ differentiation, limit DAMP signaling, or modulate IL-17A pathways may prevent amplification of neuronal injury.

In summary, our findings support a model in which PTX-induced mechanical hypersensitivity reflects context-dependent immune shaping rather than simple drug accumulation. The mLD PTX regimen promotes selective immune programming dominated by CD8^+^ T cell diversification, whereas the sHD PTX regimen induces diversity through broad CD4^+^ helper expansion. Furthermore, IL-17A and granzyme A emerge as likely mediators linking adaptive immunity to neuronal dysfunction. CIPN may therefore represent a self-sustaining neuroimmune process—analogous to autoimmune disease—underscoring the need for early, mechanism-based intervention strategies.

## Acknowledgments

We would like to thank the UNE COBRE Behavior Core, especially Dr. Tamara King and Denise Giuvelis, for assisting in experimental design, analysis, and interpretation.

## Generative AI

ChatGPT (free version, OpenAI, GPT-4o) was only used to assist with rephrasing for clarity, grammatical improvements, and flow. AI was not used for analysis, interpretation, or content. Authors are solely responsible and confirm the accuracy of content within this publication.

