## Supplemental Materials for "Paclitaxel Dosing Regimens Drive Differential CD4⁺ T Cell Responses in the Dorsal Root Ganglia and Modulate Neuropathic Pain Severity"

**Supplemental Tables**

| Antibody/Dye | Vendor, Cat # | Epitope: surface (S), intracellular (I) |
|---|---|---|
| Live/Dead Fixable Violet Dye | ThermoFisher, L34955 | NA |
| CD3 eF506 | ThermoFisher, 69-0032-82 | S |
| IL-17A BV605 | Biolegend, 506927 | I |
| CD8 BV650 | Biolegend, 100742 | S |
| IL-2 AF 488 | ThermoFisher, 53-7021-82 | I |
| LAP PerCP-eFluor™ 710 | ThermoFisher, 46-9821-82 | I |
| Granzyme A PE | ThermoFisher, 12-5831-82 | I |
| CD4 PE-Texas Red | ThermoFisher, MCD0417 | S |
| FoxP3 PerCP-Cy5.5 | ThermoFisher, 45-5773-80 | I |
| IL-4 PE-Cy7 | ThermoFisher, 25-7041-82 | I |
| Perforin APC | ThermoFisher, 17-9392-80 | I |
| IL-10 AF 700 | ThermoFisher, 56-7101-82 | I |
| IFN-γ APC-eFluor 780 | ThermoFisher, 47-7311-82 | I |

**Table S1.** Flow panel used to determine cytokine producing cells in the DRG.

| Antibody/Dye | Vendor | Epitope: surface (S), intracellular (I) |
|---|---|---|
| Live/Dead Fixable Violet Dye | ThermoFisher, L34955 | NA |
| CD3 eF506 | ThermoFisher, 69-0032-82 | S |
| NK1.1 SB600 | ThermoFisher 63-5941-80 | S |
| CD8 BV650 | Biolegend, 100742 | S |
| CD45 FITC | Biolegend, 160305 | S |
| CD11c PE-Cy5.5 | ThermoFisher 35-0114-80 | S |
| CD4 PE-Texas Red | ThermoFisher, MCD0417 | S |
| CD68 PerCP/Cyanine5.5 | Biolegend, 137010 | I |
| CD19 PE-Cyanine7 | ThermoFisher, 25-0193-82 | S |
| MHC Class II, I-A/I-E APC | ThermoFisher, 17-5321-82 | S |
| Ly-6G AF700 | Biolegend, 127622 | S |
| CD11b APC-eFluor 780 | ThermoFisher, 47-0112-82 | S |

**Table S2.** Flow panel used to determine $CD45^{+}$ immune cell populations in the DRG.

**Tables S3** and **S4** found in **Supplemental Excel File**.

### Supplemental Figures

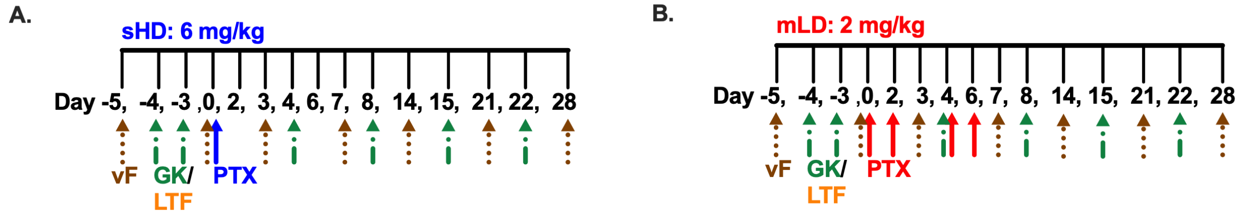


**S1 Fig. Timeline of mechanical sensitivity testing with CD4⁺ T cell depletion and PTX dosing. (A, B)** Mechanical hypersensitivity (tactile threshold, grams) was assessed using von Frey filaments (brown arrow) on day (D) -5. To deplete CD4⁺ T cells, naïve female mice received GK1.5 (CD4⁺ T cell depleting antibody) injections (green arrows) on D -4, -3. Control mice received the LTF.2 isotype control antibody (orange) on the same schedule as GK1.5. Mechanical sensitivity was reassessed on D 0 to determine whether CD4⁺ T cell depletion or isotype control treatment altered baseline sensitivity. This D 0 value was used as the baseline (BL) measurement prior to PTX administration. Two PTX dosing regimens were then used: **(A)** a single high dose of 6 mg/kg PTX administered on D 0 (sHD, blue) or **(B)** multiple low doses of 2 mg/kg PTX administered on D 0, 2, 4, 6 (mLD, red). Mechanical hypersensitivity was assessed by von Frey testing on D 3, 7, 14, 21, and 28 to evaluate the effects of each dosing regimen. To prevent re-population of CD4⁺ T cells, additional GK1.5 injections were administered on D 4, 8, 15, and 22. LTF.2 isotype control antibody injections followed the same schedule as GK1.5 injections.

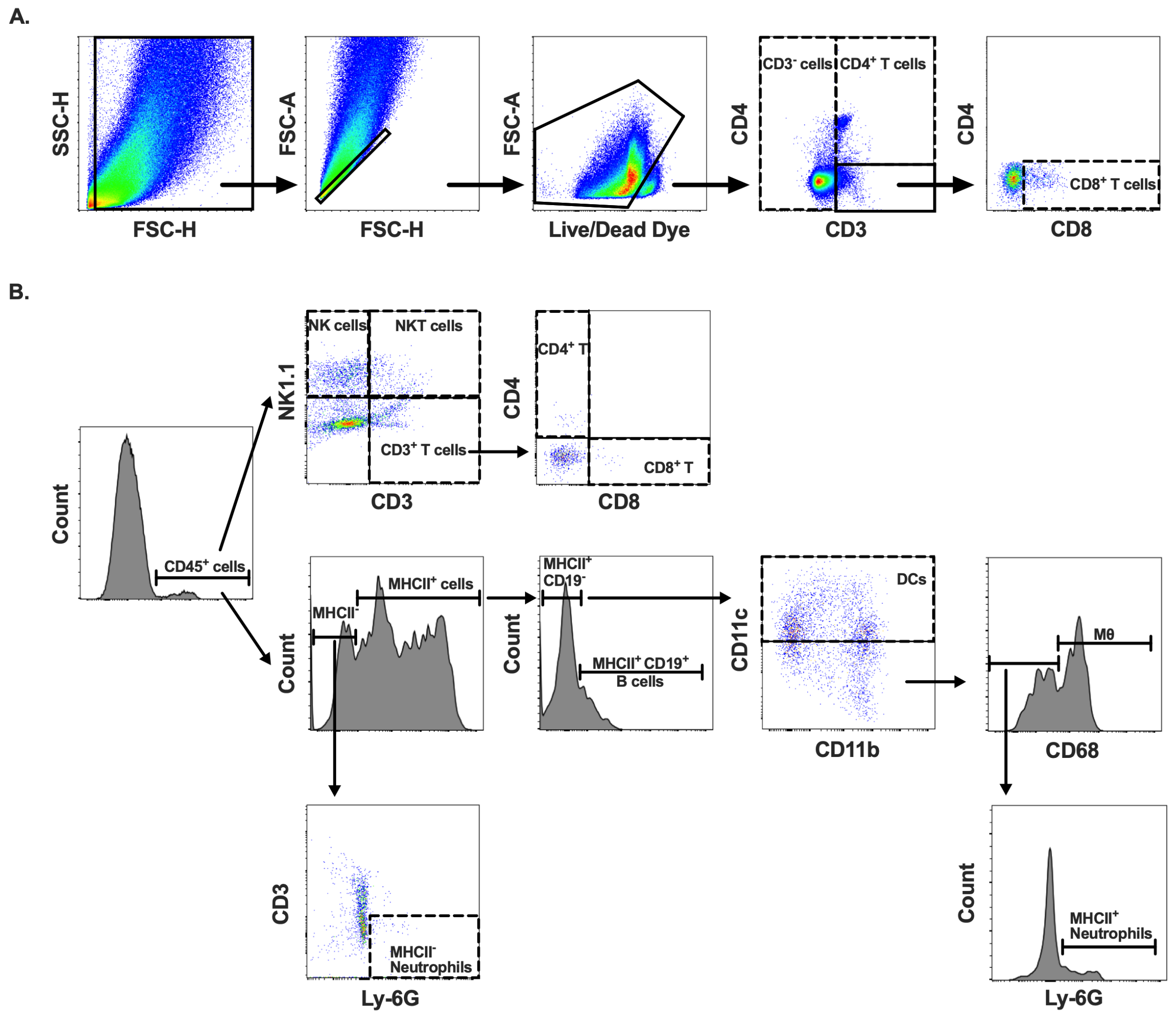


**S2 Fig. Multi-color flow cytometry gating strategies for immune cell populations in the DRG. (A)** A nested gating strategy was used to identify immune cells within the DRG. Acutely dissociated dorsal root ganglion (DRG) cells were gated based on size (forward scatter-height, FSC-H) and granularity (side scatter-height, SSC-H) (column 1). From the total cell gate (column 1), single cells were determined by plotting the cell height against the cell area. From the single cell gate (column 2), live cells were identified based on the exclusion of the viability violet dye. From the live cell gate (column 3), $CD4^+$ T cells were determined by plotting CD3 against CD4 (column 4). A gate was drawn around $CD3^+$ $CD4^-$ cells and plotted against CD8 to identify $CD8^+$ T cells (column 5). **(B)** Gating strategy for $CD45^+$ immune cell populations in the DRG.

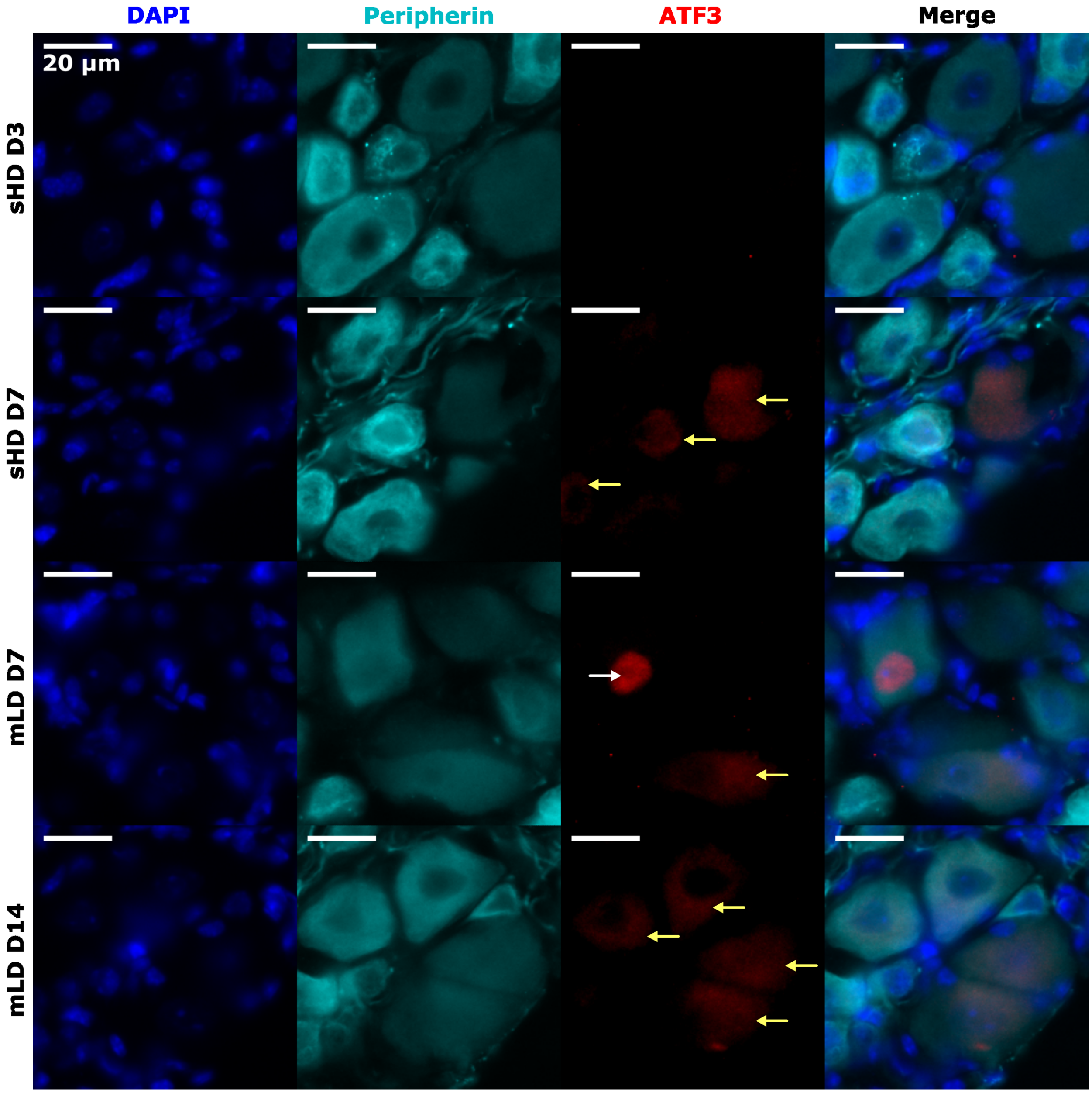


**S3 Fig. Neuronal ATF3 after sHD and mLD PTX treatment.** Representative widefield epifluorescence images of peripherin (teal) L4 DRG neurons from sHD PTX (days (D) 3, 7) and mLD PTX (days (D) 7, 14) female mice. Sub-cellular location of ATF3 (red) is indicated by arrows: yellow denotes cytoplasmic expression, and white denotes nuclear localization (co-localized with DAPI, blue).

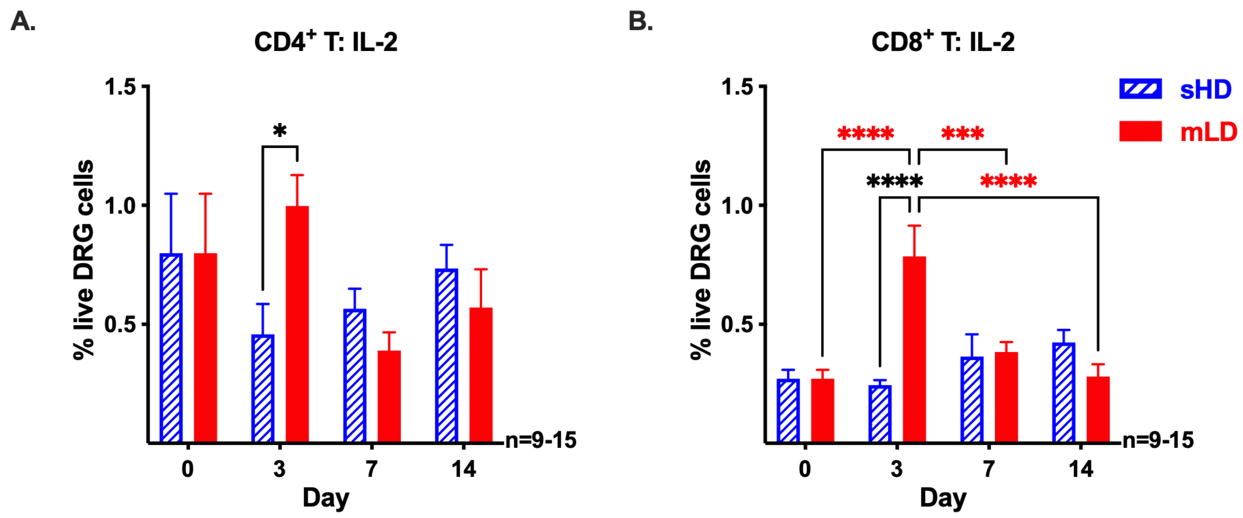


**S4 Fig. The mLD PTX regimen induces a stronger IL-2 T cell response in the DRG compared to the sHD PTX regimen.** Frequency of IL-2 producing **(A)** $CD4^+$ T cells and **(B)** $CD8^+$ T cells out of total live DRG cells on day 0 and after sHD (blue) or mLD (red) PTX regimens. Significance was determined by 2-way ANOVA with Tukey's multiple comparisons test (*p<0.05, ***p<0.001, ****p<0.0001, n=9-15/regimen). Blue (sHD) and red (mLD) significance asterisks compare T cell frequencies within each dosing regimen. Black significance asterisks compare T cell frequencies between sHD and mLD PTX regimens.

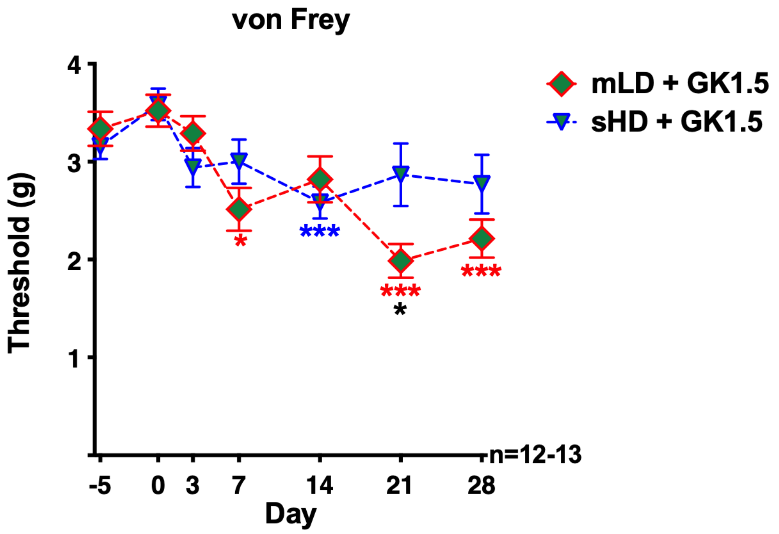


**S5 Fig. CD4$^{+}$ T cell depletion led to a greater reduction in mechanical hypersensitivity with the sHD PTX regimen compared to the mLD PTX regimen.** Mechanical hypersensitivity (tactile threshold, grams) was assessed in CD4$^{+}$ T cell deficient (GK1.5) mice with von Frey filaments prior to antibody injection (D -5), prior to PTX (D 0) to determine baseline (BL) values, and after sHD or mLD PTX on D 3, 7, 14, 21, and 28. Blue (sHD) and red (mLD) significance asterisks compare hypersensitivity after PTX to BL within each group. Black significance asterisks compare hypersensitivity values between mLD + GK1.5 and sHD + GK1.5.

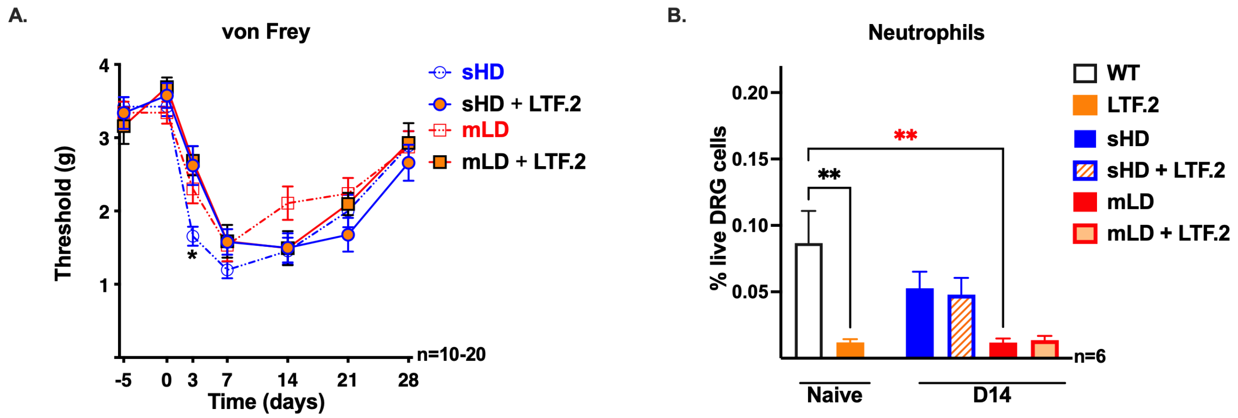


**S6 Fig. LTF.2 isotype control antibody reduces neutrophils in the DRG, and delays PTX-induced mechanical hypersensitivity in the sHD group. (A)** Comparison of PTX-induced mechanical hypersensitivity in the presence or absence of the LTF.2 isotype control antibody. Black asterisk indicate a significant difference in hypersensitivity between sHD and sHD + LTF.2 at D 3. **(B)** Frequency of neutrophils out of total live DRG cells in the presence or absence of LTF.2 isotype control antibody in naïve and D14 sHD and mLD treated mice. Significance was determined by 1-way ANOVA with Tukey's multiple comparisons test (**p<0.01, n=6). Red significance asterisk compares neutrophil cell frequencies between naïve and mLD D14. Black significance asterisks compare cell frequencies between naïve WT (absence of LTF.2) and naïve + LTF.2.

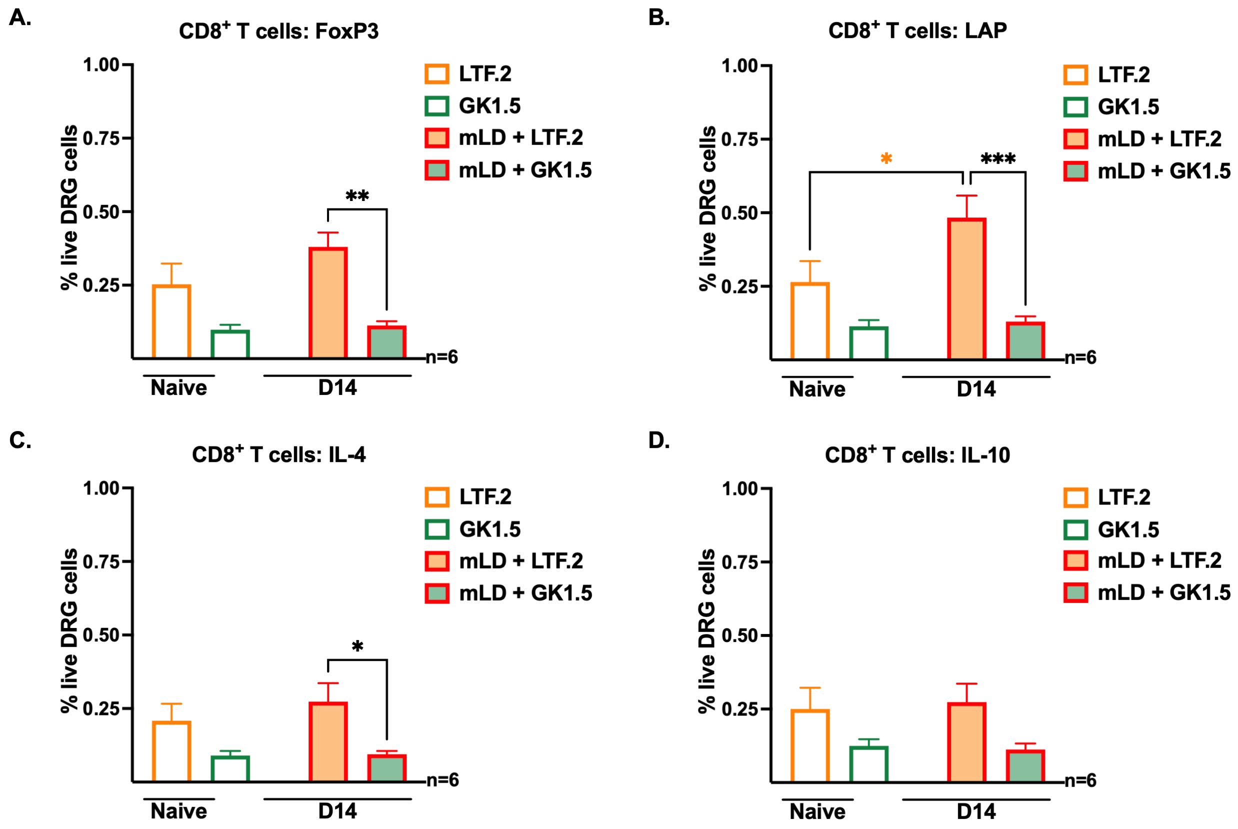


**S7 Fig. CD4⁺ T cell depletion decreases the frequencies of anti-inflammatory CD8⁺ T cells in the DRG after mLD PTX regimen.** Frequencies of **(A)** FoxP3, **(B)** LAP, **(C)** IL-4, **(D)** IL-10 CD8⁺ T cells out of total live cells in DRG from naïve mice treated with LTF.2 (white bar with orange border) or GK1.5 (white bar with green border), and from mLD D14 mice treated with LTF.2 (orange bar with red border) or GK1.5 (green bar with red border). Significance was determined by 1-way ANOVA with Tukey's multiple comparisons test (*p<0.05, **p<0.01, ***p<0.001, n=6). Orange significance asterisk compares cell frequencies between naïve and mLD D14 within LTF.2 antibody group. Black significance asterisks compare cell frequencies between LTF.2 and GK1.5 groups.
